# Reverse Engineering the Programming Logic of Cytoskeletal Dynamics

**DOI:** 10.1101/2025.09.10.674348

**Authors:** David Larios, Bibi Najma, Jiapei Miao, Shichen Liu, Heun-Jin Lee, Rob Phillips, Matt Thomson

**Author notes:** Contributing authors.

## Abstract

Eukaryotic cells generate mechanical forces through cytoskeletal filaments actively reorganized by families of molecular motors. However, how motor sequence specifies filament organization dynamics remains unclear. We developed ActiveDROPS, a cell-free platform that expresses kinesin variants in bacterial lysate droplets and quantitatively maps the resulting microtubule dynamics into a common phenotype space. Across a library of twelve kinesin-1 homologs, we observed three phenotypes: “Slow-Sustained” flows that activate after 8 h, reach peak mean velocities of 24 nm/s and decay after ***∼***32 h; “Fast-Burst” flows that activate within minutes and dissipate within 2 h, reaching velocities up to 900 nm/s; and a ***∼***36-h “Multiphase” progression through nematic, rotational, and contractile states. Microtubule gliding assays and molecular dynamics simulations using AlphaFold-predicted structures linked the “Fast-Burst” phenotype to generally faster motility and more favorable motor–tubulin interactions than “Slow-Sustained”. By replacing the microtubule-binding region of a “Slow-Sustained” motor with that of a “Fast-Burst” motor, we generated a “Fast-Sustained” chimera with flows that activate within 1 h, peak at 70 nm/s and persist for 16 h, showing that recombination can reprogram the parental relationship between speed and timing of microtubule-motor self-organization. These results reveal a constrained logic through which kinesin sequence shapes cytoskeletal dynamics, providing a framework for dissecting the mechanical repertoire available to cellular systems.

## 1 Introduction

Living cells continually reshape their internal architecture to divide, move, transport material, and respond to the environment. These functions are accomplished by generating mechanical force through networks of protein filaments actively organized by molecular motors (Ronceray et al., 2016; Soares e Silva et al., 2011; Xie and Minc, 2020). Despite vast phenotypic diversity across eukaryotes, the high-level architecture of this cytoskeletal system remains remarkably conserved. The main components of the cytoskeleton, actin filaments and microtubules, assemble from small, energy-hydrolyzing subunits and are dynamically regulated by a broad set of accessory proteins and motors (Brouhard and Rice, 2014; Goodson and Jonasson, 2018; Manka and Moores, 2018). These networks are not passive scaffolds; they are active matter: materials that remodel continuously and support behaviors as diverse as mitotic spindle formation and cell migration (Carvalho et al., 2013; Fürthauer and Shelley, 2022; Sanchez et al., 2012). Since early in evolution, eukaryotes evolved a genetic toolbox of cytoskeletal proteins with a remarkably high number of kinesin motor variants, each with different kinetic properties tailored to build large-scale organized structures such as the spindle, cilia, or intracellular transport networks (Belsham et al., 2022; Hirokawa et al., 2009; Scarabelli et al., 2015; Vale and Fletterick, 1997; Yue et al., 2018). However, how cells harness the individual properties of motor variants (“choose tools from the toolbox”) and accessory proteins to assemble complex mechanical machinery remains poorly understood.

The basic building block of cytoskeletal function is motor-filament self-organization (Lam et al., 2016; Nédélec et al., 1997; Surrey et al., 2001). It has been extensively shown that motors and filaments alone in ATP solution self-organize into dynamical phases, from turbulent flows and vortices that resemble cytoplasmic streaming, or contractile asters like those that form during cell division (Foster et al., 2015; Gordon et al., 2012; Head et al., 2011; Henkin et al., 2014; Kruse et al., 2004; Sanchez et al., 2012). These phases can be accessed simply by changing the relative amounts of motors to filaments, and can be “steered” towards other dynamical states by associated proteins or crosslinking factors (Najma et al., 2024; Roostalu et al., 2018; Stanhope et al., 2017). This suggests that cells can already access a range of self-organizing behavior by tuning the expression of motors in its genetic toolbox, and perform mechanical “operations” steering self organization with its multiple layers of regulation to exert control over cytoskeletal filament dynamics in space and time (Bidone et al., 2017; Fürthauer et al., 2019; Striebel et al., 2020). To understand the logic by which these cytoskeletal components assemble into large-scale mechanical machinery, we first need to identify the core “operations” accessible through the sequence space of motor proteins.

However, in vitro reconstitution provides controlled access to network mechanics, but purification is laborious and often requires construct-specific optimization, making broad sequence comparisons difficult (Jijumon et al., 2024; Laohakunakorn et al., 2020; Liu and Fletcher, 2009; Noireaux and Liu, 2020). In vivo knockouts and perturbations can reveal the cellular functions of individual proteins, but the assembly process of intracellular mechanical machinery remains obscured by cellular complexity (Dammann and Köster, 2014; Liu and Fletcher, 2009; Vogel et al., 2013). Here we introduce ActiveDROPS, a high-throughput approach to quantitatively map kinesin motor genetic sequences to microtubule self-organization dynamics. Libraries of kinesin motor genes are individually mixed into E. coli lysate (Garamella et al., 2016) droplets containing pre-stabilized microtubule networks, generating different spatiotemporal patterns of self-organization that are determined by the biochemical features encoded in the sequence of each motor. By imaging multiple droplets in parallel, we quantitatively map these resulting dynamics to a low-dimensional phenotype space, allowing the systematic exploration of phenotypic diversity in genomic databases. Moreover, these phenotypes can be mechanistically dissected by modifying the input DNA and remapping it to phenotype space, thus learning the rules by which structural features propagate up to macroscopic self-organization.

Across twelve kinesin-1 homologs, ActiveDROPS revealed two recurring mechanical phenotypes: Slow-Sustained flows with delayed activation and prolonged activity, and Fast-Burst flows with rapid activation and decay. A divergent motor, ThTr, generated a Multiphase phenotype progressing through nematic organization, coherent rotation, and global contraction. Microtubule-gliding assays and molecular dynamics simulations linked Fast-Burst dynamics to generally faster ATP-dependent motility and more favorable computed motor–tubulin interactions than Slow-Sustained dynamics. We then used recombination to test how these contrasting collective behaviors depend on motor architecture. Replacing the region containing the microtubule-binding interface of a Slow-Sustained motor with the corresponding region from a Fast-Burst motor generated a distinct Fast-Sustained phenotype, combining faster network motion than the Slow-Sustained parent with longer activity than the Fast-Burst parent. Across the recombinant library, the outcome of each exchange depended on the surrounding sequence, showing that these collective behaviors arise from compatible combinations of coupled motor regions. Finally, varying ThTr expression through DNA titration shifted whether networks remained nematic, reached rotation, or completed the Multiphase progression through contraction. Together, these results demonstrate that coupled elements of motor architecture encode the spatiotemporal trajectory of macroscopic self-organization and provide a framework for dissecting the mechanical repertoire available to cellular systems

## 2 Results

### 2.1 ActiveDROPS maps kinesin motor sequences to emergent microtubule network dynamics

We developed ActiveDROPS (Active Dynamic Reprogrammable self-Organizing Protein System) to explore and dissect how sequence-level variation in kinesin motor genes determine microtubule organization dynamics. Each reaction combines a linear DNA template encoding a kinesin–mVenus fusion (Fig. S1), bacterial cell-free transcription–translation, and a preassembled microtubule network within a water-in-oil droplet (Fig. 1A). The droplet format isolates individual reactions in stable, small-volume compartments that permit oxygen flow, allowing multiple motor variants and replicate droplets to be arrayed across multi-well plates and followed for tens of hours. Automated widefield fluorescence microscopy repeatedly images multiple droplets in parallel, recording Cy5-labeled microtubules together with kinesin–mVenus accumulation, estimated from a fluorescence calibration series (Fig. S2). As the encoded motor accumulates, ATP-driven motor–microtubule interactions collectively activate filament motion and reorganize the initially disordered network (Fig. 1B,C). Expression and self-organization thus unfold within the same compartment, enabling high-throughput measurements that link a programmable DNA input directly to the network dynamics it generates without purifying each motor in advance.

**Fig. 1.**
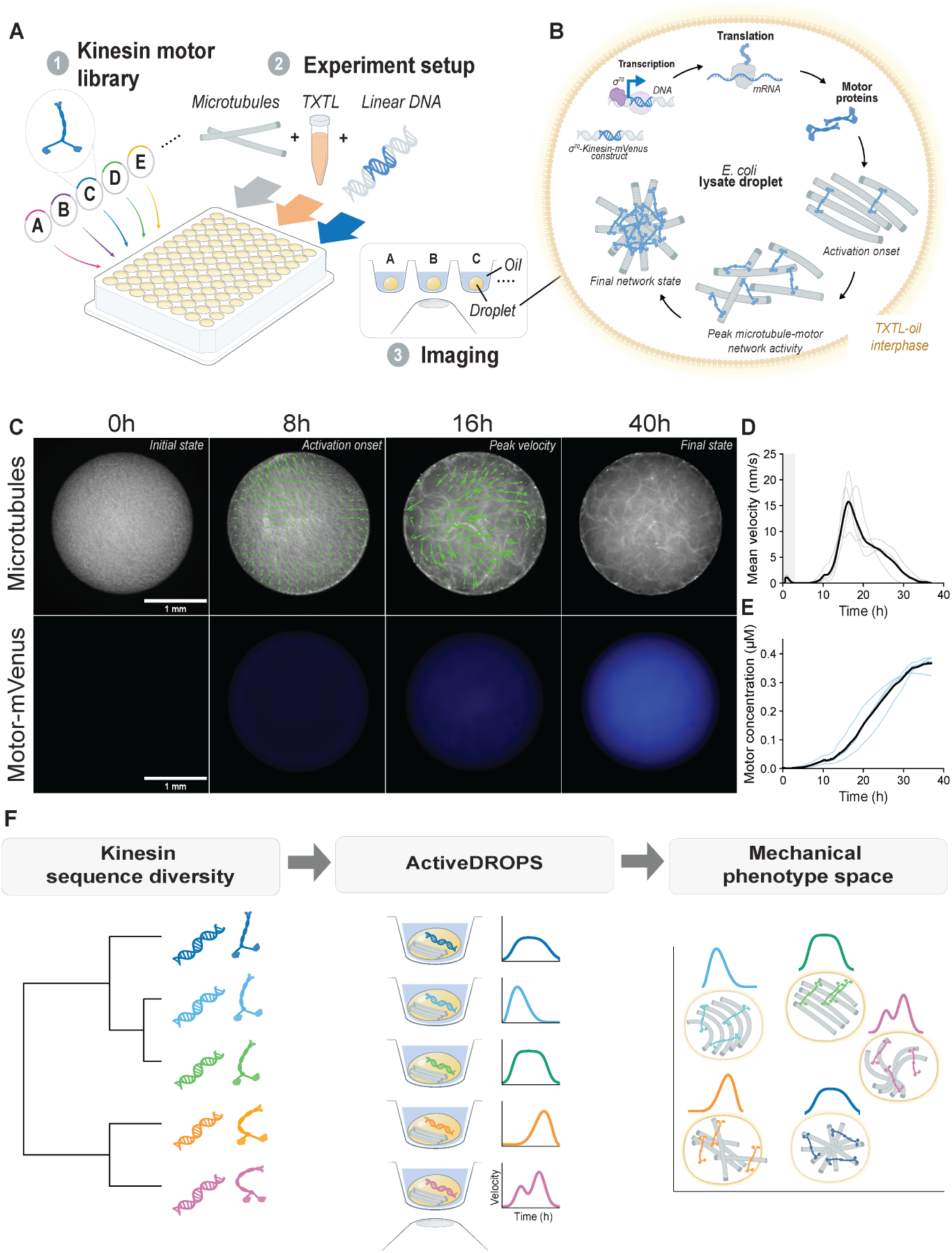
ActiveDROPS reconstitutes microtubule-motor self-organization from kinesin gene expression. (A) Experimental framework. Linear DNA templates encoding kinesin-1 motors are combined with pre-polymerized, GMP-CPP–stabilized microtubules and E. coli TXTL lysate, then compartmentalized into water-in-oil droplets for dual-channel time-lapse imaging. (B) Schematic of coupled gene expression and cytoskeletal dynamics within a single droplet. Motors are transcribed and translated from DNA templates and engage the microtubule network as they accumulate. (C) Representative time-lapse widefield epifluorescence microscopy of a droplet expressing Drosophila K401 (100 nM DNA). Top row: microtubule network (Cy5, grayscale) with PIV-derived velocity fields (green arrows). Bottom row: motor protein fluorescence (mVenus). The initially disordered network transitions into coherent flows as motor concentration rises (Supplementary Video 1). Scale bar, 1000 *µ*m. (D) Mean microtubule flow velocity over time for K401 at 100 nM DNA, showing activation at approximately 10 h and peak velocity of approximately 16 nm/s at approximately 16 h. (E) Motor protein concentration over time, measured by mVenus fluorescence calibrated to absolute molarity. (F) Conceptual overview of the ActiveDROPS workflow mapping kinesin sequence diversity to mechanical phenotype space.

We utilized the model kinesin motor K401, a well-characterized Drosophila melanogaster kinesin-1 construct, as a workhorse to develop our platform. Calibrated mVenus fluorescence estimated the accumulation of the K401–mVenus reporter, while particle image velocimetry (PIV) applied to successive images of Cy5-labeled microtubules generated velocity fields and the droplet-averaged network velocity (Fig. 1C–E; Fig. S2). Across four independently prepared droplets, network velocity remained near baseline for approximately 10 h, rose to a pointwise mean maximum of approximately 16 nm/s near 16 h, and then declined over the remainder of the experiment. Individual droplets developed distinct spatial flow fields, yet preserved a common temporal program of activation, duration, and velocity, showing that K401 expression generates a reproducible mechanical phenotype despite the stochastic nature of microtubule-motor self-organization (Fig. 1D,E).

Because parallel imaging produces many long time-lapse movies, we use a quantitative framework to compare the complete dynamics of every droplet. For each droplet, we retain the mean network velocity over absolute experimental time, preserving both the magnitude and duration of activity. We then use nonnegative matrix factorization (NMF) to identify recurring temporal velocity motifs across the trajectories. Each droplet is represented by how strongly these motifs contribute to its trajectory, placing droplets with similar dynamics near one another in a low-dimensional mechanical phenotype space (Fig. 1F). The number of motifs is selected by cross-validation based on reconstruction of held-out portions of the velocity trajectories. We then identify groups of droplets by clustering their complete set of standardized motif weights. Motor identity, mVenus reporter concentration, and qualitative phenotype labels are excluded from both model selection and clustering. The resulting groups are compared with the qualitative phenotypes observed in the movies only after clustering. Two-dimensional projections are used only for visualization. This framework allows mechanical phenotypes to emerge from the data while retaining differences in when motion begins, how fast the network moves, and how long its activity persists.

### 2.2 Sequence variation maps onto three distinct mechanical phenotypes

Previous studies have shown that kinesin-1 motors retain a conserved motor structure while differing in nucleotide dependence, ATP turnover, processivity, and regulation across evolutionary lineages (Steinberg and Schliwa, 1996; Kirchner et al., 1999; Kallipolitou et al., 2001; Röhlk et al., 2008). These observations suggest that sequence divergence within kinesin-1 can encode substantial biochemical diversity without disrupting the overall motor structure. To test whether this diversity also propagates into collective mechanical behavior, we used K401 and Kif3 as well-characterized animal and amoebozoan reference motors and assembled a library of ten additional uncharacterized kinesin-1 homologs from animals, amoebozoans, and other unicellular eukaryotes (Fig. 2A; Table S1). Pairwise sequence identity across the aligned motor regions ranged from 35.8% to 91.4% and was generally higher within the animal and amoebozoan groups than between groups. In contrast, structural identity (symmetric average-length-normalized US-align scores) for the corresponding AlphaFold-predicted structures ranged from 0.794 to 0.959, revealing high structural similarity across the library despite substantial sequence divergence (Fig. 2B). We used ActiveDROPS to ask whether sequence or structural similarity predicts mechanical phenotype, and whether the biochemical diversity previously observed among individual kinesins propagates into distinct collective dynamics.

**Fig. 2.**
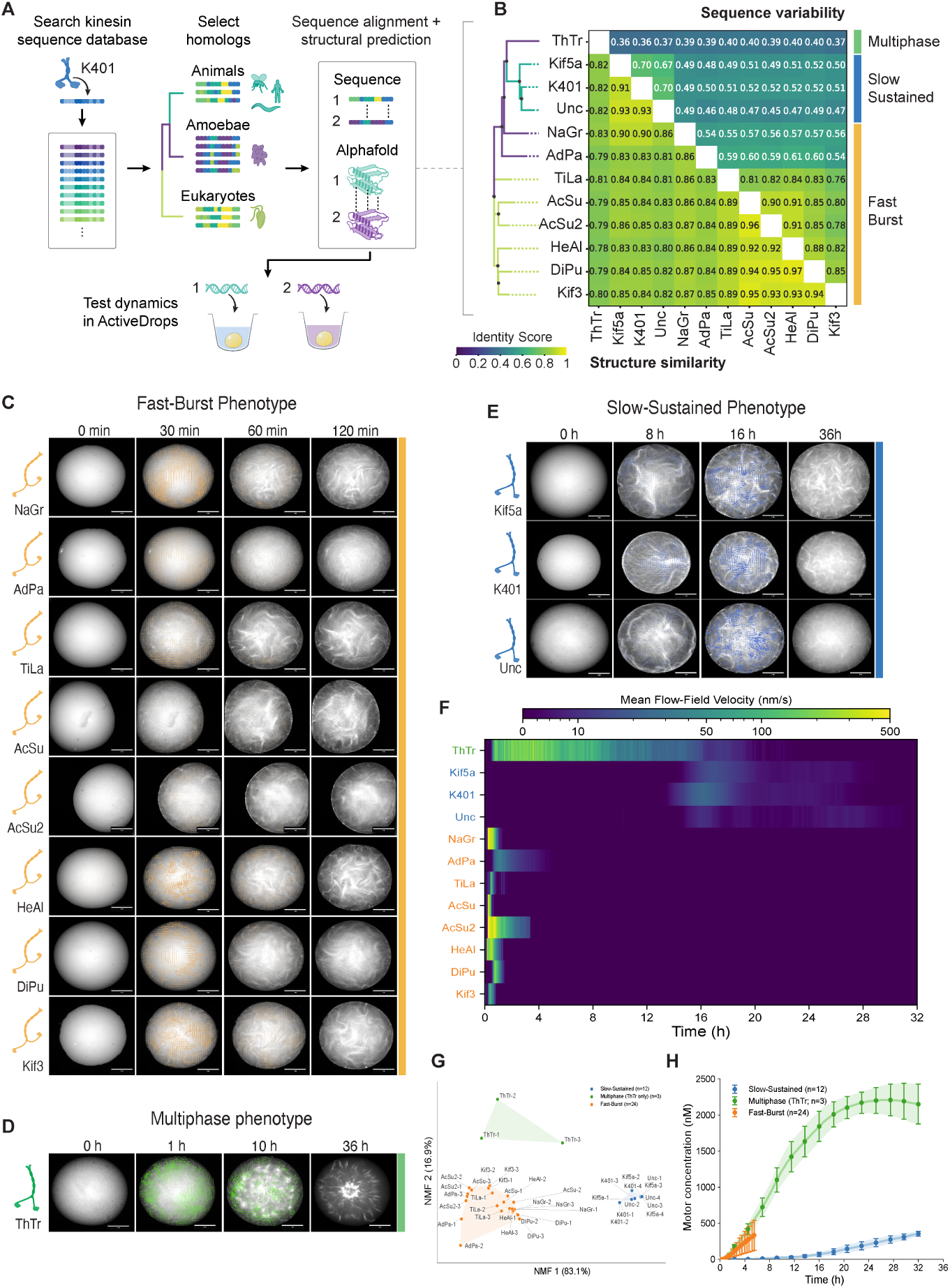
A motor protein library generates two conserved and one divergent mechanical phenotypes. (A) Selection of twelve kinesin-related sequences for structural and mechanical comparison. (B) Pairwise sequence identity (upper triangle) and symmetric average-length-normalized US-align score (lower triangle). The displayed order is fixed rather than inferred from the similarity matrix. (C) Representative Fast-Burst trajectories from eight motors (Supplementary Video 2a). (D) Representative ThTr progression from nematic organization through rotation to contraction. (E) Representative Slow-Sustained trajectories from Kif5a, K401, and Unc (Supplementary Video 2b). Supplementary Video 2c compares all three patterns. Scale bars in C–E, 1 mm. (F) Mean PIV velocity for one representative preparation per motor. Recorded Fast-Burst intervals after confirmed cessation of motion are displayed as zero; unrecorded intervals are left missing. (G) Velocity-only NMF phenotype space independently fitted to all 39 droplets. Cross-validation selected three temporal components, and label-free clustering in the complete standardized NMF-weight space identified three clusters. Colors denote the resulting clusters, and droplet identities and phenotype annotations were added only after fitting. The axes show the first two principal components of the three-dimensional NMF-weight space and are used only for visualization; clustering was performed before this two-dimensional projection. (H) Estimated motor concentration from calibrated kinesin–mVenus fluorescence, shown as preparation-level mean *±* SD for Slow-Sustained (n = 12), Fast-Burst (n = 24), and Multiphase (n = 3) droplets.

Despite their high structural similarity, the twelve motors generated two conserved and one divergent mechanical phenotypes. The three animal motors—Kif5a, K401, and Unc—produced a Slow-Sustained phenotype. Across replicates, these motors reached a median processed peak velocity of 11.8 nm/s at 16.8 h, and the median final recorded time above 20% of peak velocity was 31.7 h. In contrast, eight motors produced a Fast-Burst phenotype, reaching a median processed peak velocity of 59.3 nm/s at 0.33 h and a median final recorded time above 20% of peak velocity of 1.33 h across replicate droplets. The Fast-Burst phenotype included amoebozoan motors as well as NaGr and AdPa from other unicellular eukaryotes, despite their 54–59% sequence identity with members of the amoebozoan group. The most sequence-divergent motor, ThTr from Thecamonas trahens, produced a Multiphase phenotype. During its accumulation trajectory, ThTr progressed through nematic and turbulent organization, droplet-scale rotation, and a final global contraction. Slow-Sustained and Fast-Burst therefore differed primarily in their velocity and temporal profiles, while both generated stochastic turbulent flows. ThTr instead accessed additional forms of spatial organization.

For each motor, we measured at least three independently prepared droplets. Although individual droplets produced different flows, all replicate droplets expressing the same motor grouped together in the velocity-only NMF phenotype space, and the label-free analysis recovered the Slow-Sustained, Fast-Burst, and Multiphase phenotypes (Fig. 2G). Motor identity was also associated with distinct accumulation profiles: at 4 h, estimated motor concentrations were 3.1 *±* 2.4 nM for Slow-Sustained motors, 236.4 *±* 152.6 nM for Fast-Burst motors, and 271.7 *±* 90.9 nM for ThTr (mean *±* SD; Fig. 2H). These results connect motor sequence to both accumulation kinetics and collective dynamics under common bacterial-lysate conditions. We next used microtubule-gliding assays to test whether the collective phenotypes were also associated with differences in motor function.

### 2.3 Microtubule gliding assays and molecular dynamics simulations reveal the biochemical basis of the two conserved mechanical phenotypes

Earlier studies established that conventional animal kinesins and fast amoebal/fungal-type kinesins occupy distinct kinetic regimes. In microtubule-gliding assays, in which surface-immobilized motors propel fluorescent, stabilized microtubules across a coverslip, amoebal and fungal motors generally produced several-fold higher gliding velocities than animal kinesins and required higher ATP concentrations to reach half-maximal velocity (Saxton et al., 1988; Steinberg and Schliwa, 1996; Kirchner et al., 1999; Röhlk et al., 2008). The recurrence of Slow-Sustained and Fast-Burst phenotypes among motors from these lineages raised the possibility that the two conserved mechanical phenotypes reflect corresponding differences in motor mechanochemistry. We therefore measured ATP-dependent microtubule gliding for each of the twelve motors under common assay conditions (Fig. 3A,B; Fig. S7). Gliding velocities were fit to the Michaelis–Menten relationship, (*v*([ATP]) = *V*_max_[ATP]*/*(*K*_half_,_ATP_ + [ATP])), yielding the saturating gliding velocity, *V*_max_, and the apparent ATP concentration required to reach half-maximal velocity, *K*_half_,_ATP_. Dividing *V*_max_ by the 8-nm kinesin step size provided an apparent stepping-rate estimate and the corresponding ratio of apparent stepping rate to *K*_half_,_ATP_.

**Fig. 3.**
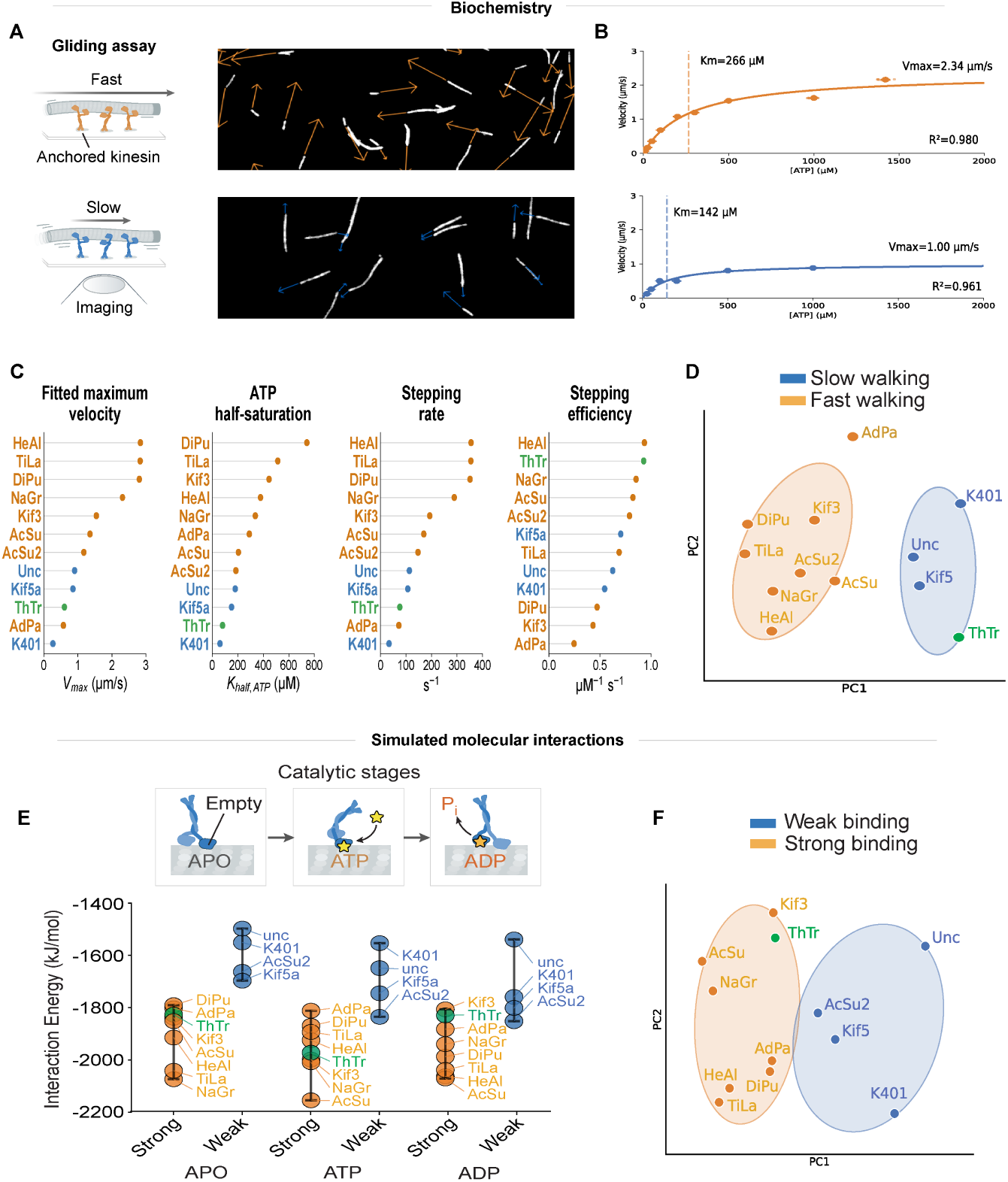
Microtubule gliding assays and molecular dynamics simulations reveal the biochemical basis of the two conserved self-organization patterns. (A) Microtubule gliding assay. Surface-immobilized kinesin motors propel fluorescent microtubules across a coverslip; representative Fast-Burst (orange) and Slow-Sustained (blue) trajectories are shown (Supplementary Video 3). (B) Representative Michaelis–Menten fits for HeAl (orange; *V*_max_ = 2.84 *µ*m s*^−^*^1^, *K*_half_,_ATP_ = 379 *µ*M) and Kif5a (blue; *V*_max_ = 0.86 *µ*m s*^−^*^1^, *K*_half_,_ATP_ = 152 *µ*M). Dashed lines mark the fitted apparent *K*_half_,_ATP_. (C) Ranked kinetic descriptors for the twelve motors: saturating gliding velocity, apparent *K*_half_,_ATP_, apparent stepping rate calculated using an 8-nm step size, and the ratio of apparent stepping rate to *K*_half_,_ATP_. (D) Principal component analysis of the standardized kinetic descriptors. (E) Short-range Coulombic plus Lennard-Jones motor–tubulin interaction energies in apo, ATP-bound, and ADP-bound simulations. Points and error bars show the mean and standard deviation across three trajectories per motor and state; more negative values indicate more favorable direct interactions and are not binding free energies. (F) Principal component analysis of the three nucleotide-state interaction-energy profiles. Orange, blue, and green denote Fast-Burst, Slow-Sustained, and Multiphase motors, respectively.

Fast-Burst and Slow-Sustained motors occupied largely distinct kinetic ranges. The eight Fast-Burst motors reached *V*_max_ values of 0.57–2.84 *µ*m s*^−^*^1^, corresponding to apparent stepping rates of 72–355 s*^−^*^1^. In contrast, the three Slow-Sustained motors reached *V*_max_ values of 0.27–0.90 *µ*m s*^−^*^1^ and apparent stepping rates of 33–113 s*^−^*^1^. AdPa overlapped the Slow-Sustained velocity range, with the lowest *V*_max_ among the Fast-Burst motors despite its Fast-Burst class assignment. The classes also differed in their ATP dependence: Slow-Sustained motors reached half-maximal velocity at 61– 181 *µ*M ATP, whereas Fast-Burst motors required 186–742 *µ*M ATP. ThTr combined a comparatively low *V*_max_ of 0.60 *µ*m s*^−^*^1^ with a low apparent *K*_half_,_ATP_ of 81 *µ*M. The fitted parameters and their derived quantities therefore broadly separated the classes of microtubule organization while retaining AdPa and ThTr as informative exceptions (Fig. 3B–D; Table S2).

Because force generation also depends on how kinesin interacts with the microtubule lattice, we next used atomistic molecular dynamics simulations to examine whether the mechanical phenotypes were associated with differences at the motor– tubulin interface. Each of the twelve motor heads was modeled in complex with the same human *αβ*-tubulin heterodimer using AlphaFold3, allowing sequence-dependent interactions to be compared against a common tubulin background. We simulated each complex in nucleotide-free apo, ATP-bound, and ADP-bound states to compare the interfaces under distinct nucleotide conditions relevant to motor stepping. Three independent 50-ns trajectories were generated for each motor and nucleotide state. We then calculated the mean short-range Coulombic and Lennard-Jones interaction energy between motor and tubulin over the final 30 ns of each trajectory (Fig. 3E). More negative values indicate more favorable direct nonbonded interactions under matched simulation conditions, providing a comparative molecular descriptor of how each motor interacts with tubulin rather than a measurement of binding affinity.

The molecular dynamics simulations revealed two broad interaction-energy ranges across the three nucleotide states. The Slow-Sustained motors K401, Kif5a, and Unc occupied the less favorable range, whereas seven of the eight Fast-Burst motors occupied the more favorable range; AcSu2 lay closer to the Slow-Sustained group. Principal component analysis of the three state-specific energies retained this broad separation (Fig. 3F; Table S3). ThTr combined favorable interaction energies comparable to Fast-Burst motors with relatively slow gliding, distinguishing its molecular profile from the two recurring phenotypes. Together, the gliding assays and simulations associate Fast-Burst organization with generally faster motility and more favorable motor–tubulin interactions, and Slow-Sustained organization with slower motility and less favorable interactions. AdPa and AcSu2 depart from the prevailing kinetic and interaction-energy trends, respectively, while ThTr produces Multiphase organization. ATP-dependent motility and motor–tubulin interactions therefore provide complementary molecular coordinates linking kinesin sequence to collective dynamics, raising the question of how these properties are coupled within the motor architecture.

### 2.4 Reconfiguring motor protein architecture rewrites the self-organization dynamics of microtubule networks

The biochemical and computational characterization of the motor library showed that the Slow-Sustained and Fast-Burst mechanical phenotypes occupy distinct mechanochemical regimes, broadly separated by ATP-dependent gliding kinetics and simulated motor–tubulin interaction energies across the nucleotide cycle. Motor stepping depends on coordinated conformational changes linking nucleotide processing, microtubule binding, and force transmission. We therefore asked whether sequence differences between K401 (Slow-Sustained) and Kif3 (Fast-Burst) could transfer specific features of their mechanical phenotypes, or whether the effect of each region depended on the surrounding motor architecture. Guided by established kinesin structure–function relationships, aligned parental sequences, and AlphaFold-predicted structures, we divided the motors into three operational crossover regions (Fig. 4A,B). M1 spans the nucleotide-binding core and contains the P-loop, *α*0 helix, Loop 5, and central *β*-sheet that organize the ATP-binding pocket (Behnke-Parks et al., 2010; Endow et al., 2010; Wang et al., 2015). M2 contains Switch I and Switch II together with Loop 11, *α*4, *α*5, and Loop 12, linking nucleotide-state transitions to the principal microtubule-binding interface (Atherton et al., 2014; Brier et al., 2006; Nitta et al., 2004; Sindelar and Downing, 2007). M3 contains the neck linker and proximal coiled coil involved in transmitting conformational changes from the motor head, directed stepping, and dimerization (Romberg et al., 1998; Skiniotis et al., 2003; Tripet et al., 1997). The structural elements and mechanochemical roles of the three operational regions are summarized in Table S4. We then generated the complete 2^3^ library containing the two parental motors and all six recombinant configurations (Fig. 4C). The S and F designations indicate whether each region was derived from K401 or Kif3, respectively, and do not assign an intrinsic slow, fast, sustained, or burst property to any individual region.

**Fig. 4.**
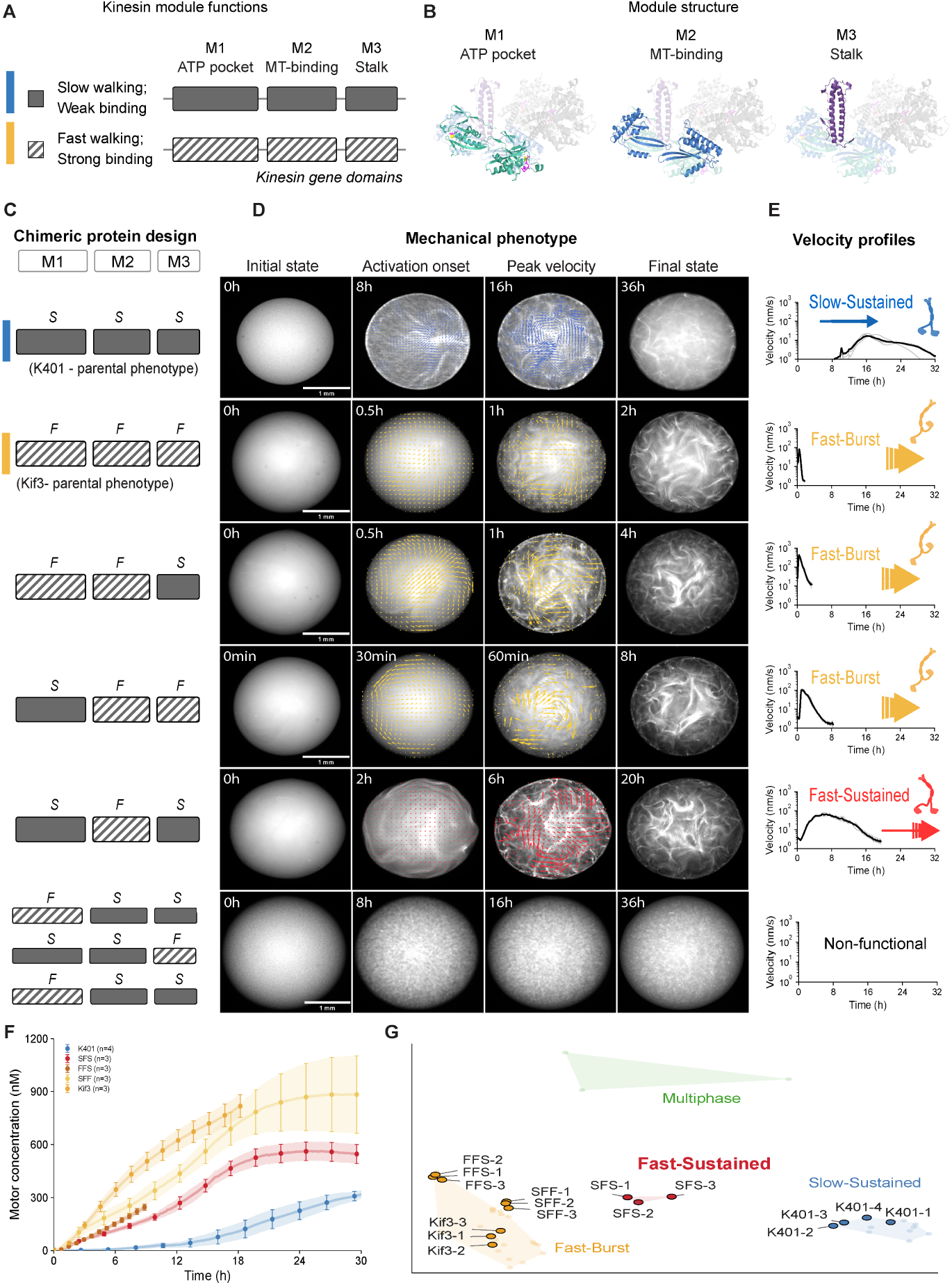
Reconfiguring motor protein architecture rewrites the self-organization dynamics of microtubule networks. (A) Operational M1–M3 crossover scheme. M1, M2, and M3 span the ATP-processing pocket, microtubule-binding interface, and proximal coiled coil, respectively. The regions contain distributed, coupled functions rather than independent modules. S and F denote K401- and Kif3-derived sequences. (B) Locations of the three regions in the predicted motor–tubulin structure. (C) Complete 2^3^ parental and recombinant design. (D) Representative time-lapse images for K401, Kif3, and the three motile recombinants (Supplementary Video 4). The lower row contains visually immobile designs for which comparable PIV trajectories were unavailable. Scale bars, 1 mm. (E) Velocity profiles for the parental and motile recombinant motors. (F) Estimated motor concentration from calibrated kinesin–mVenus fluorescence, shown as mean *±* SD across independent preparations (K401, n = 4; each other displayed construct, n = 3). (G) Velocity-only NMF phenotype space for 48 natural-motor and motile-chimera trajectories. Label-free clustering was performed in the complete NMF-weight space; the first two principal components are shown for visualization. SFS formed a distinct cluster containing no parental motor or other active chimera.

Using K401 (SSS) as the starting architecture, we first replaced each region independently. Replacing M1 alone (FSS) or M3 alone (SSF) produced no detectable collective motion under the tested conditions. In contrast, replacing only M2 generated an active SFS motor with dynamics distinct from either parent. Relative to K401, SFS increased the mean peak velocity from 18.8 to 71.0 nm/s and shifted the mean peak time from 16.72 to 6.22 h, while retaining prolonged activity: the mean durations above half-maximum were 7.85 h for K401 and 6.74 h for SFS. SFS therefore combined faster network motion than K401 with longer activity than Kif3, reconfiguring the parental speed–duration relationship. Within the K401-derived M1/M3 background, the Kif3-derived M2 region generated a distinct Fast-Sustained phenotype.

SFS then provided a functional mixed architecture in which to examine additional substitutions. Replacing its K401-derived M1 with the corresponding Kif3 region to generate FFS shifted the mean peak time from 6.22 to 0.37 h and shortened the mean half-maximal duration from 6.74 to 0.60 h, producing Fast-Burst dynamics. Replacing M3 instead to generate SFF also shortened the activity period, reducing the mean half-maximal duration to 1.48 h. Across the complete library, every configuration containing the Kif3-derived M2 region—SFS, FFS, SFF, and FFF—generated detectable collective motion. Among configurations containing the K401-derived M2 region, only the parental SSS motor was active; the mixed FSS, SSF, and FSF configurations produced no detectable motion. Separate expression titrations detected reporter accumulation for FSS and FSF, arguing against a simple absence of fusion-protein production, although these measurements cannot determine whether the motors were correctly folded, oligomerized, or mechanically engaged. State-dependent motor–tubulin interaction energies and M2 interface contacts across the recombinant library are shown in Fig. S8. Full-length production of all eight chimeric constructs was assessed by anti-mVenus western blotting (Fig. S9). The library therefore identifies operational compatibility between regions while showing that the phenotype produced by a substitution depends on the surrounding sequence architecture.

At the same DNA input, all three active chimeras accumulated kinesin–mVenus earlier than K401 and reached higher estimated motor concentrations during the first 8 h (Fig. 4F). Despite this shared pattern of earlier accumulation, the chimeras generated distinct mechanical outcomes: FFS and SFF produced Fast-Burst dynamics, whereas SFS sustained network motion for hours. The NMF phenotype space recovered this distinction quantitatively, grouping all three independently prepared SFS droplets together in a cluster containing no parental motor or other active chimera (Fig. 4G). SFS therefore opened access to a distinct Fast-Sustained region of mechanical phenotype space that was not populated by either parent or the other motile chimeras. Recombining compatible regions of the motor architecture thus expanded the collective dynamics accessible from the two parental motors.

### 2.5 A divergent motor accesses three different regions of phenotype space

The motor library and recombinant chimeras showed that sequence and architecture constrain the velocity and time dimensions of microtubule self-organization. We next asked how expression moves a fixed motor within these constraints. Across classes, DNA titration produced distinct effects on motor expression rate and work (Fig. S10A,B). For a Slow-Sustained motor, reducing DNA lowered velocity and power, reducing work—the integral of power over time—while activation and decay occurred at nearly the same times (Fig. S10C–E). For a Fast-Burst motor, reducing DNA delayed and broadened the burst; lower velocity was offset by longer duration, leaving work broadly similar across inputs (Fig. S10F–H). Expression therefore scaled the magnitude of Slow-Sustained motion but redistributed Fast-Burst motion in time. ThTr presented a qualitatively different possibility: this divergent motor generated an ordered progression through nematic, rotational, and contractile organization. We therefore tested whether expression could select among spatial phases rather than only rescale motion within a turbulent-flow phenotype.

We varied ThTr expression through a stepwise tenfold titration of input DNA. At 1, 10, and 100 nM DNA, the network remained nematic, progressed into coherent rotation, or completed the nematic–rotational–contractile sequence, respectively (Fig. 5A–C). These outcomes were accompanied by increasing motor concentration and expression rate: at 12 h, estimated ThTr–mVenus concentrations were 420, 749, and 1,159 nM, and maximum expression rates were 58, 88, and 140 nM/h for the one, two, and three phase outcomes (Fig. 5D,F). Peak velocities nevertheless remained comparable at 416, 408, and 384 nm/s, showing that progression through additional phases did not arise from faster motion (Fig. 5E). Instead, the mean activity endpoint shifted from approximately 33 h to 18 h and 15 h after the start of the experiment as the network advanced through the sequence. Velocity correlation length further distinguished persistent nematic organization, coherent rotation, and contraction (Fig. 5G). Expression therefore selected where the network terminated along a spatial phase progression encoded by ThTr.

**Fig. 5.**
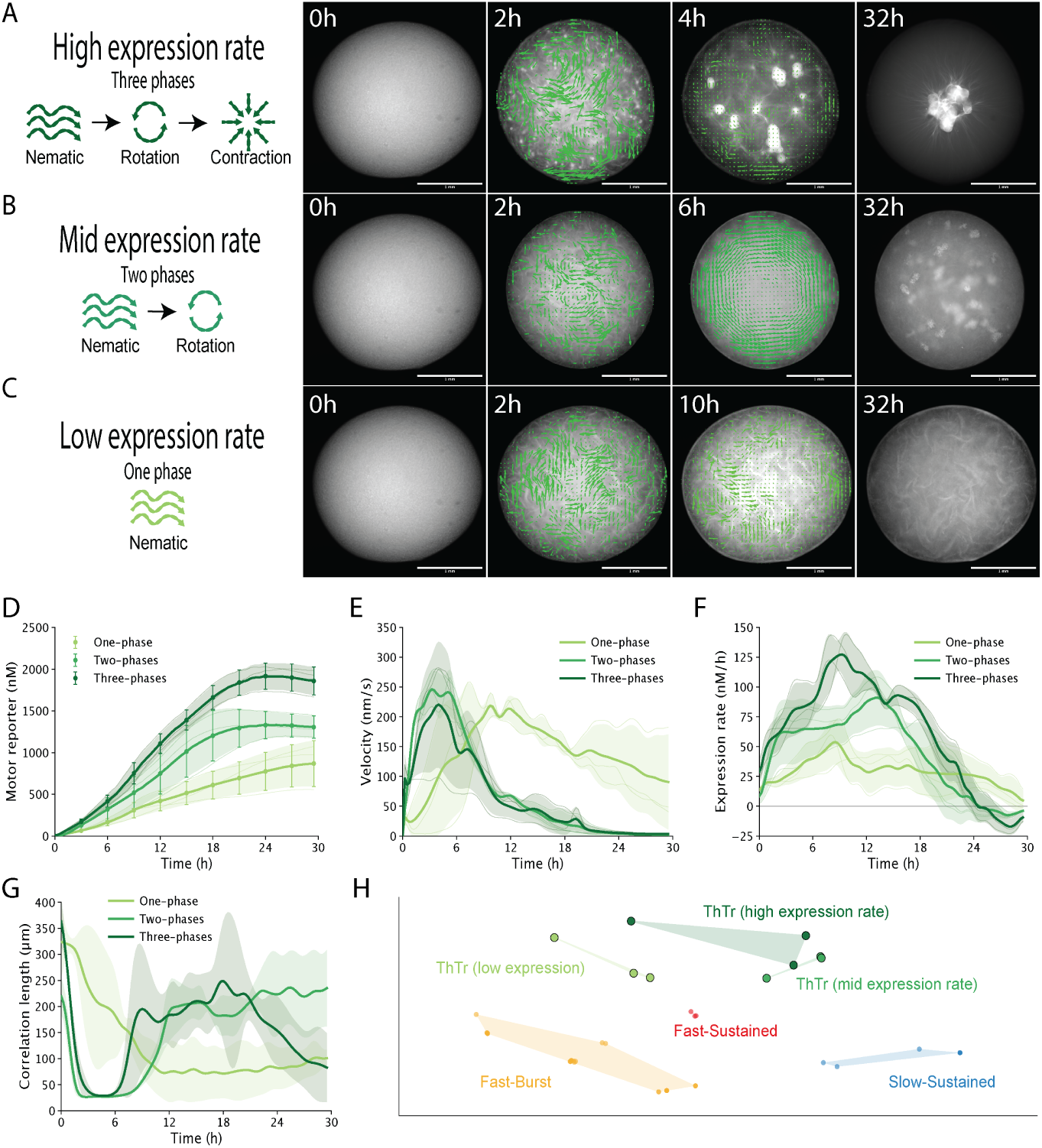
Expression control selects distinct endpoints in a divergent three-phase network progression. (A–C) Representative ThTr-driven networks under conditions producing high, intermediate, or low reporter accumulation. The networks progressed through three phases (nematic– rotation–contraction), two phases (nematic–rotation), or remained nematic, respectively. Green vectors show PIV-derived velocity fields. Scale bars, 1 mm. (D) Estimated ThTr–mVenus concentration. (E) Mean network velocity. (F) Net appearance rate of fluorescent ThTr–mVenus, not direct translation rate. (G) Velocity correlation length. Curves show individual preparations and mean *±* SD (n = 3 per outcome). (H) Velocity-only NMF behavior space fitted without motor, expression, concentration, or phase labels. The one-phase trajectories formed a distinct cluster, whereas the two- and three-phase trajectories shared a cluster. Axes show a two-dimensional visualization of the complete NMF-weight space; shaded regions are descriptive envelopes. Supplementary Video 5 shows the complete three-phase progression.

Together, the natural-motor library, recombinant chimeras, and expression titrations generated six observable mechanical outcomes: Slow-Sustained, Fast-Burst, Fast-Sustained, and the nematic, rotational, and contractile outcomes of ThTr (Fig. 5H). The space was constructed from network mechanics without using motor identity, DNA input, motor concentration, or phase labels. Rotational and contractile ThTr trajectories remained close in the velocity-only projection because they reached similar speeds, but their spatial organization and correlation-length dynamics distinguished their endpoints. Together, these experiments identify three ways to access distinct regions of phenotype space. Selecting different motor sequences accesses different dynamical regimes; recombining coupled regions of motor architecture generates new velocity–duration relationships; and tuning expression moves a fixed sequence within its accessible repertoire. For ThTr, this last operation selects among qualitatively different spatial phases. Motor sequence and architecture therefore determine the dynamics available to a microtubule network, while motor expression determines which of those dynamics is realized.

## 3 Discussion

Kinesin-microtubule interactions are elementary operations from which cells construct larger cytoskeletal structures and flows. Motors convert ATP hydrolysis into directed motion and force, but neither velocity nor binding affinity alone specifies what a population of motors will make a filament network do over time. Such collective organization contributes to intracellular transport, cell division, migration, and cytoplasmic remodeling (Ronceray et al., 2016; Soares e Silva et al., 2011; Xie and Minc, 2020). Here, we use mechanical phenotype in an explicitly restricted sense. An organismal phenotype emerges from the interaction of genotype with developmental and environmental context; the phenotypes measured here instead describe the reproducible onset, magnitude, duration, progression, and endpoint of self-organization generated by a motor sequence within a defined reconstructed environment. This operational definition isolates the elementary behaviors that a motor makes available to a microtubule network before additional cellular regulation is imposed. It therefore provides a tractable connection between sequence variation and the raw mechanical repertoire from which cytoskeletal machinery can be assembled (Brouhard and Rice, 2014; Goodson and Jonasson, 2018; Lam et al., 2016; Manka and Moores, 2018; Nédélec et al., 1997; Surrey et al., 2001).

Reconstituted motor-filament systems have demonstrated that relatively few components can generate turbulent flows, vortices, asters, contraction, and other organized states (Foster et al., 2015; Gordon et al., 2012; Head et al., 2011; Henkin et al., 2014; Kruse et al., 2004; Sanchez et al., 2012). These behaviors can be redirected by changing motor concentration, filament concentration, crosslinking, or associated proteins (Najma et al., 2024; Roostalu et al., 2018; Stanhope et al., 2017). The biological objective of the present work was to add motor sequence as an experimentally accessible coordinate in this phase space. To recover that relationship, we connected library construction, cell-free expression, time-resolved imaging, and quantitative phenotype mapping into a common workflow. A compact measurement, the mean network velocity over absolute experimental time, retained sufficient information for label-free NMF to recover Slow-Sustained, Fast-Burst, and Multiphase trajectories without using motor identity or expression labels. This representation provides a scalable way to compare coding-sequence libraries while preserving the timing and magnitude of collective activity. It also creates a basis for iterative mutate-screen-map experiments, addressing limitations of protein-purification workflows that make broad sequence comparisons difficult (Jijumon et al., 2024; Laohakunakorn et al., 2020; Liu and Fletcher, 2009; Noireaux and Liu, 2020).

Across the natural-motor library, sequence identity and predicted structural similarity did not recover the observed mechanical phenotypes. NaGr, for example, shared less than 57% sequence identity with every other motor tested but generated the same Fast-Burst trajectory as several more closely related homologs. Gliding assays and computed motor-tubulin interaction energies nevertheless identified molecular correlates of the two recurring phenotypes: Fast-Burst motors generally moved faster and formed more favorable simulated interactions than Slow-Sustained motors. This relationship is consistent with established differences in kinesin kinetics and ATP dependence across evolutionary lineages (Kallipolitou et al., 2001; Kirchner et al., 1999; Röhlk et al., 2008; Saxton et al., 1988; Steinberg and Schliwa, 1996). However, exceptions showed that no individual molecular coordinate determines the collective trajectory. Kinesin motion instead depends on coupling among nucleotide processing, microtubule binding, force transmission, and neck-linker dynamics (Atherton et al., 2014; Wang et al., 2015). The chimeras directly exposed this context dependence. Domain exchanges that previously altered single-motor velocity or processivity (Adio et al., 2009; Kalchishkova and Böhm, 2008; Shastry and Hancock, 2010) here rewrote the temporal dynamics of an entire network. The Fast-Sustained SFS chimera therefore demonstrates that recombination can open a region of mechanical phenotype space occupied by neither parent, without implying that these combinations were naturally selected or adaptive.

ThTr revealed a second layer of biological control that was not captured by the conserved mechanochemical coordinates measured here. Increasing its expression moved the network through persistent nematic organization, coherent rotation, and global contraction, while peak velocities remained comparable across the three endpoints. Expression therefore selected among a nested repertoire of states rather than merely increasing the speed of a fixed behavior. Related transitions in reconstituted active networks can arise from motor accumulation, confinement, filament alignment, and competition between extensile and contractile interactions (Doostmohammadi et al., 2018; Juniper et al., 2018; Lemma et al., 2022; Najma et al., 2024; Sumino et al., 2012; Suzuki et al., 2017). ThTr suggests that an apparently complex progression of network states need not require a comparably complex sequence of regulatory interventions. A single motor can make multiple behaviors accessible, with abundance determining where the progression terminates. The responsible molecular features remain unknown, but plausible candidates include motor-motor association, oligomeric state, cooperative microtubule binding, and force-dependent attachment. Distinguishing among these mechanisms will require measurements of active-motor fraction, oligomerization, residence time, and load-dependent kinetics.

The bacterial lysate provides a shared environment for comparing sequences, but it is not biologically neutral. It lacks native eukaryotic chaperones, post-translational modifications, interaction partners, and degradation pathways, and mVenus fluorescence reports accumulated fusion protein rather than the mechanically active fraction. The slower accumulation of the animal motors may consequently reflect differences in synthesis, maturation, folding, solubility, or stability rather than an intrinsic taxonomic distinction. Earlier reporter accumulation was compatible with distinct mechanical outcomes, as the active chimeras generated either Fast-Burst or Fast-Sustained dynamics. Cell-free synthesis has previously produced functional kinesins in gliding assays (Inoue et al., 2023), supporting its use as a bridge from sequence to motor activity while emphasizing the importance of expression context. Future comparisons using eukaryotic lysates, dynamic microtubules, mixed motors, and microtubule-associated proteins can determine how this logic changes as cellular regulation is restored (Dammann and Köster, 2014; Garamella et al., 2016; Vogel et al., 2013). Together, the results define a programming logic in which motor sequence constrains the accessible mechanical repertoire, coupling across motor architecture configures its spatiotemporal trajectory, and expression selects the network state ultimately realized.

## 4 Conclusion

Kinesin sequence variation encodes a constrained repertoire of microtubule self-organization dynamics. This repertoire is not specified through independent sequence modules or captured by a single biochemical property. Instead, it emerges from distributed coupling among nucleotide processing, microtubule binding, and force transmission across the motor architecture. Recombination can reconfigure this coupling and generate velocity-duration relationships absent from the parental motors, while expression determines which states within a sequence-dependent repertoire are ultimately reached. These findings define a biological programming logic in which motor sequence establishes the possibilities for collective behavior and regulation selects among them. By connecting molecular variation to macroscopic organization, this work provides a route to understanding how evolution can diversify cytoskeletal dynamics while retaining a conserved motor architecture.

## Materials and methods

### Kinesin constructs and linear DNA templates

Each expression template contained the p70a *σ*70 promoter and a ribosome-binding site followed by a kinesin motor region comprising the catalytic motor domain, neck linker, and a proximal coiled coil selected to preserve dimerization. The motor sequence was fused through a flexible (GS)_4_ linker to mVenus, followed by an AviTag, a 6*×*His tag, and the T500 terminator (Fig. S1). The same expression architecture was used for the twelve natural kinesin homologs, the K401 and Kif3 parental constructs, and the six recombinant constructs. Coding sequences were codon-optimized for expression in *Escherichia coli*, obtained as synthetic gene fragments, and assembled into the common expression cassette. Linear templates were amplified with a high-fidelity DNA polymerase using primers flanking the complete expression cassette, purified by silica-column cleanup, and quantified by absorbance and fluorescence-based double-stranded-DNA measurements. DNA inputs and the relationship between preparations and figures are retained in the accompanying dataset and replicate inventory. Motor identities, source sequences, and truncation boundaries are reported in Table S1.

### Design and sequence verification of the K401–Kif3 recombinant library

The K401 and Kif3 sequences were aligned and divided into three operational crossover regions using established kinesin structure–function relationships and the corresponding AlphaFold-predicted structures. M1 contains the nucleotide-binding core, including the P-loop, alpha-0 helix, Loop 5, and central beta-sheet that organize the ATP-binding pocket. M2 contains Switch I and Switch II together with Loop 11, alpha-4, alpha-5, and Loop 12, linking nucleotide-state transitions to the principal microtubule-binding interface. M3 contains the neck linker and proximal coiled coil involved in transmitting conformational changes from the motor head, directed stepping, and dimerization. The crossover coordinates were residues 1–179, 180–334, and 335–401 for K401 and residues 1–175, 176–330, and 331–592 for Kif3.

All eight combinations of the three regions were constructed. Each construct was named according to the parental origin of M1, M2, and M3, with S denoting sequence from K401 and F denoting sequence from Kif3. These letters identify sequence origin only and do not assign a slow, fast, sustained, or burst property to an individual region. Benchling GenPept exports were treated as the authoritative experimental sequence records. Each A–H construct was reconstructed computationally from its three annotated parental segments and compared residue by residue with the corresponding experimental sequence and with chain A of the structure used for simulation. All eight sequences matched the expected parental-segment combinations with zero residue mismatches.

### Microtubule preparation

GMP-CPP-stabilized microtubules were prepared in a 50-*µ*L polymerization reaction containing 80 mM K-PIPES, pH 6.8, 1 mM EGTA, 2 mM MgCl2, 75 *µ*M unlabeled tubulin, 5 *µ*M Cy5-labeled tubulin (PurSolutions), 1 mM DTT, and 0.6 mM GMP-CPP. Before polymerization, the mixture was centrifuged at approximately 300,000 *× g* for 5 min at 2 *^◦^*C to remove aggregates. The supernatant was transferred to a fresh tube and incubated at 37 *^◦^*C for 1 h. Filament lengths were measured from surface-immobilized preparations; the resulting population had a mean length of 16.7 *µ*m (Fig. S3).

### ActiveDROPS reactions and droplet formation

Each 4.5-*µ*L reaction was prepared by combining 2.5 *µ*L of the myTXTL Linear DNA Expression master mix (Arbor Biosciences), 1.0 *µ*L of GMP-CPP-stabilized microtubules, and 1.0 *µ*L of linear DNA solution. The completed reaction contained 11 *µ*M total tubulin and 100 nM linear DNA, except for the ThTr DNA-dose experiment in Results section 2.5, which used final DNA concentrations of 1, 10, or 100 nM. Glass-bottom 96-well plates were passivated overnight with 0.5% BSA in PBS, rinsed with PBS followed by water, air-dried, and filled with 300 *µ*L of light mineral oil per well. The mineral oil contained no phospholipid. A 1.0-*µ*L aliquot of the completed reaction was manually deposited into the oil in each well to form a single aqueous droplet at the glass surface.

### Time-lapse microscopy

ActiveDROPS droplets were imaged on a Nikon Ti2-E inverted microscope equipped with a Plan Apochromat 4*×*, NA 0.20 objective, a Lumencor SOLA SE light source, Cy5 and YFP/mVenus filter sets, and a Hamamatsu ORCA-Flash 4.0 sCMOS camera. Acquisition was controlled with Micro-Manager 2.0. Fluorescent microtubules and kinesin–mVenus were recorded in separate channels with 2 *×* 2 camera binning. The acquisition interval ranged from 6 to 60 s depending on the duration and temporal resolution required for each experiment. Original timestamps, spatial calibration, exposures, and image-derived measurements were retained in the source tables. These data were acquired by widefield epifluorescence microscopy; the term confocal is not used for this imaging configuration.

### mVenus calibration and estimation of motor accumulation

Purified concentrated mVenus standards spanning 0–320 ng *µ*L*^−^*^1^ were encapsulated and imaged using the same droplet geometry and optical settings used for ActiveDROPS measurements (Fig. S2). Mean droplet fluorescence was linear across the calibration series and was fit by

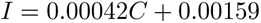

where *I* is the background-corrected fluorescence intensity and *C* is the mVenus concentration in micrograms per milliliter. Standard concentrations were converted to molarity using the molecular mass of mVenus. Because every motor polypeptide contained one C-terminal mVenus, the calibrated signal was reported as the estimated concentration of accumulated kinesin–mVenus fusion protein. This measurement does not separately determine fluorophore maturation, protein degradation, protein folding, oligomerization, or the fraction of motor that is mechanically active.

### Particle image velocimetry and flow-field measurements

Microtubule motion was quantified in MATLAB with PIVlab v3.0 using two-pass FFT cross-correlation with interrogation windows of 180 pixels in the first pass and 90 pixels in the second pass, with 50% overlap. A circular mask excluded pixels outside the droplet. At each time point, the mean flow-field velocity was calculated as the mean magnitude of the valid velocity vectors within the mask. Where indicated, the same velocity fields were used to calculate vorticity, positive and negative divergence, strain, shear, and the spatial velocity correlation length. The correlation length was defined as the distance at which the normalized spatial velocity autocorrelation decayed to 1/*e*. Kinesin–mVenus fluorescence was measured from the corresponding reporter channel and converted to estimated motor concentration using the calibration above.

### Sequence identity and predicted-structure similarity

The motor sequences were aligned with Clustal Omega v1.2.4. The motor-domain interval defined by K401 was propagated through the alignment and used for every pairwise comparison. Pairwise sequence identity was calculated as the fraction of exact amino-acid matches among alignment columns occupied in both sequences. The corresponding segment was trimmed from each AlphaFold-predicted structure and compared with US-align. The displayed structure-similarity value was the TM-score normalized by the average length of the two trimmed structures, making the reported score symmetric with respect to comparison direction. The twelve motors produced 66 unique pairwise comparisons. Motor order in the matrix was fixed for display and was not inferred from the similarity values.

### AlphaFold structure prediction and complex preparation

Predicted structures of the truncated kinesin constructs, construct dimers, and kinesin–tubulin complexes were generated with AlphaFold3 from the corresponding protein sequences. The same default-ranked AlphaFold3 model was retained consistently for each prediction and archived with the project data. The motor–tubulin models contained a kinesin motor head bound to the same *αβ*-tubulin template so that interaction measurements could be compared across motors. The nucleotide pocket was prepared in apo, ATP-bound, or ADP-bound form for the molecular-dynamics simulations described below.

### Velocity-only NMF behavior spaces

Non-negative matrix factorization (NMF) was used to determine whether the mechanical trajectories reproduced the phenotype groupings without using motor identity or phenotype labels. The analyses associated with Figures 2, 4, and 5 were fitted independently. Consequently, the temporal components and cluster coordinates in one figure are not assumed to be identical to those in another.

For each analysis, mean-velocity histories were resampled onto an absolute-time grid with 20-min spacing, with interpolation restricted to the measured portion of each trajectory. Shorter histories were extended with zeros to the figure-specific endpoint. For Fast-Burst preparations, imaging ended after collective motion had ceased, so this extension represented continuation of the observed inactive state. The common time axis allowed brief bursts and sustained activity to be compared over the same elapsed experimental time while retaining their differences in activity duration. Appended zeros were included in the NMF input. Time was not rescaled, and individual trajectories were not normalized to their own maxima. Velocity was transformed with log1p and divided by the pooled 99.5th percentile of the transformed values in the corresponding dataset.

NMF ranks from 1 to 8 were compared by four-fold cross-validation using contiguous held-out blocks comprising approximately 15% of the local time grid per fold. Rank was chosen by the one-standard-error rule, selecting the lowest-rank model whose reconstruction error was within one standard error of the minimum. Cross-validation used six initializations per fit and a maximum of 1,200 iterations. The selected model was then refitted to the complete dataset with 12 initializations, a maximum of 1,600 iterations, and random seed 42. Component stability was evaluated across eight additional single-start fits by cosine matching of temporal components.

Droplets were clustered in the complete standardized NMF-weight space using agglomerative clustering with average linkage. Candidate cluster numbers from 2 to 8 were evaluated by silhouette score and by stability over 250 resamples containing 80% of the droplets. When solutions were otherwise comparable, preference was given to a solution in which every cluster contained at least three droplets. Principal-component analysis of the NMF weights was used only to display the first two axes; clustering was not performed in the two-dimensional projection. Motor identity, recombinant identity, estimated motor concentration, DNA input, and phenotype annotations were withheld from NMF fitting, rank selection, and clustering and were added only after the representation was fixed.

The velocity-only analyses selected three temporal components and three clusters for the 39 Figure 2 trajectories, four components and five clusters for the 48 Figure 4 trajectories, and three components and five clusters for the 54 Figure 5 trajectories. The common endpoints of these three fits were 39, 39, and 41.33 h, respectively. For the Supplementary Information method comparison, hidden Markov models were fit to the same velocity-only inputs. The number of states was selected by held-out-droplet likelihood, and complete decoded paths were compared using Hamming distance. This comparison was used to evaluate analysis robustness and did not replace the NMF representation used for the main figures.

### Microtubule-gliding assays

Gliding flow cells were assembled from an acid-cleaned coverslip and slide separated by double-sided tape. Surfaces were incubated sequentially for 5 min each with anti-His antibody diluted 1:50 in M2B, 1% Pluronic F-127 in M2B, and the unpurified TXTL reaction containing kinesin–mVenus–6*×*His. After unbound material was washed away, fluorescent GMP-CPP-stabilized microtubules were introduced at a final total tubulin concentration of 0.5 *µ*M in motility buffer containing M2B, 0.5 mg/mL casein, 1 mM MgCl2, 10 mM DTT, 1 mg/mL glucose oxidase, 0.1 mg/mL catalase, 25 mM glucose, and ATP at concentrations from 6 to 3,000 *µ*M. Movies were acquired on a Nikon Ti2 microscope with a 60*×*, NA 1.49 oil-immersion objective at 1 frame per second for 2–5 min.

### Molecular-dynamics simulations of natural motor–tubulin complexes

AlphaFold-predicted kinesin–tubulin complexes were prepared in apo, ATP-bound, and ADP-bound nucleotide states and simulated for 50 ns. The natural-motor dataset contained three independent trajectories for each of the twelve motors in each nucleotide state, for 108 trajectories in total. Simulations were performed in GROMACS 2024 with the CHARMM36m force field and explicit TIP3P water. Each complex was placed in a solvated dodecahedral box, neutralized, and brought to 150 mM NaCl. After steepest-descent minimization, systems were equilibrated for 100 ps in the NVT ensemble and 100 ps in the NPT ensemble at 300 K and 1 bar. Production trajectories were run for 50 ns with a 2-fs time step, LINCS bond constraints, particle-mesh Ewald electrostatics, and a 1.2-nm short-range cutoff.

Motor–tubulin interaction energy was calculated as the sum of the short-range Coulombic and Lennard-Jones interaction terms between the motor and tubulin. For the natural-motor comparison, each trajectory was averaged over 20–50 ns and the three replicate values were summarized for each motor and nucleotide state. These are comparative force-field interaction-energy terms. They are not enthalpies, binding affinities, or binding free energies.

Cross-scale comparisons were performed at the motor level. ThTr was excluded when comparing the two recurrent Slow-Sustained and Fast-Burst groups. State-specific and state-averaged energy summaries were standardized where required by the displayed multivariate analysis. Associations with gliding parameters were evaluated using motor-level Spearman correlations and treated as exploratory.

### Statistical analysis and reporting

Independent TXTL preparations, independently prepared ActiveDROPS droplets, independently prepared gliding video fields, and independent simulation trajectories were treated as distinct sampling units as specified above. Summary curves and scalar measurements are reported as mean *±* SD unless stated otherwise. Individual preparations are shown where space permits. Descriptive labels were applied after unsupervised fitting and were not used to train the NMF representations. No missing experimental trajectory was converted to zero except for the explicitly defined common-endpoint construction used for NMF or the stated Figure 1 initialization convention. Sample sizes, exclusions, source provenance, and the uncertainty represented by each analysis are recorded in the figure captions and Supplementary Tables.

## Author contributions

All authors contributed equally to this work.

## Supplementary Information

### Kinesin constructs and linear DNA templates

All kinesins were expressed from a common linear DNA architecture containing upstream expression-control elements followed by the kinesin-coding sequence, mVenus, an AviTag, and a C-terminal 6*×*His tag (Fig. S1). Each kinesin construct included the motor domain, neck linker, and proximal coiled-coil region selected to preserve dimerization. The motor sequence was connected to mVenus through a flexible (GS)_4_ linker. Coding sequences were codon-optimized for E. coli, synthesized as gene fragments, and assembled within the common expression cassette. Linear templates were amplified using primers outside the expression cassette, purified, and quantified before addition to the cell-free reactions.

### Calibration of mVenus and estimation of motor-concentration

Purified mVenus standards spanning 0–320 ng/*µ*l were encapsulated in water-in-oil droplets and imaged using the same droplet geometry and optical settings as the ActiveDROPS experiments (Fig. S2). Mean droplet fluorescence increased linearly with mVenus concentration over the measured range and was described by (I = 0.00042C + 0.00159), where (I) is mean fluorescence intensity and (C) is the mVenus concentration in *µ*g/ml. This calibration was used to convert kinesin–mVenus fluorescence into an estimated molar concentration trajectory for each droplet. Because the measurement reports fluorescent kinesin–mVenus fusion, it does not separately resolve fluorophore maturation, protein folding, degradation, oligomerization, or the fraction of mechanically active motor.

We asked whether the collective-dynamics phenotypes observed in the microscopy movies could be recovered from complete droplet trajectories without supplying motor identities, construct identities, expression conditions, or phenotype labels. Each droplet was represented on an absolute-time axis, with shorter recordings extended with zeros to a common endpoint within each independently analyzed figure. For Fast-Burst preparations, recording ended after motion had ceased; the extension therefore represented continued inactivity and preserved the contrast between brief and sustained activity on the common time axis. We compared NMF and Gaussian HMM representations using either mean velocity magnitude alone or seven transformed mechanical quantities. NMF rank and HMM state number were selected by cross-validation, while droplet cluster number was selected from silhouette separation and subsampling stability in the complete fitted representation. For Figures 2, 4, and 5, velocity-only NMF selected ranks of 3, 4, and 3 and identified 3, 5, and 5 clusters, respectively; the corresponding multichannel models selected ranks of 3, 3, and 4 and identified 4, 3, and 5 clusters. In Figure 2, velocity-only NMF recovered the slow-sustained, fast-burst, and multiphase phenotypes exactly when the movie-derived labels were revealed post hoc (adjusted Rand index, ARI = 1.000), as did velocity-only HMM (ARI = 1.000). In contrast, the multichannel NMF and HMM representations each introduced a fourth cluster containing a single droplet, without improving recovery of the three major phenotypes (ARI = 0.937 and 0.995, respectively). Thus, complete velocity histories are sufficient to resolve the principal collective-dynamics phenotypes without manually constructed trajectory summaries or supervised labels. NMF provides the more interpretable primary representation because its components are explicit temporal motifs and its droplet weights quantify their relative contributions, while HMM supplies an independent comparison based on complete hidden-state paths (Figs. S4–S6).

---

**Fig. S1.**
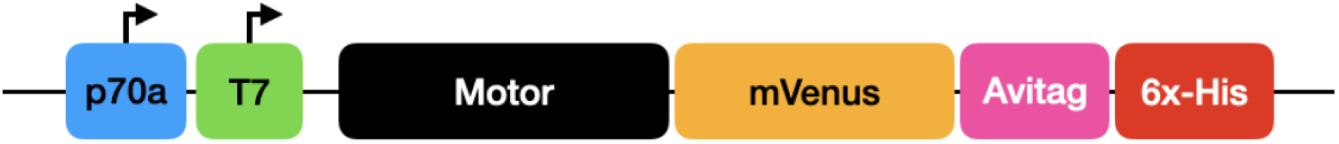
Linear DNA architecture used for kinesin expression. Schematic of the common expression template containing the upstream expression-control region, kinesin motor sequence, mVenus reporter, AviTag, and C-terminal 6*×*His tag. The motor-coding sequence was varied while the surrounding expression architecture was maintained across the kinesin library.

**Fig. S2.**
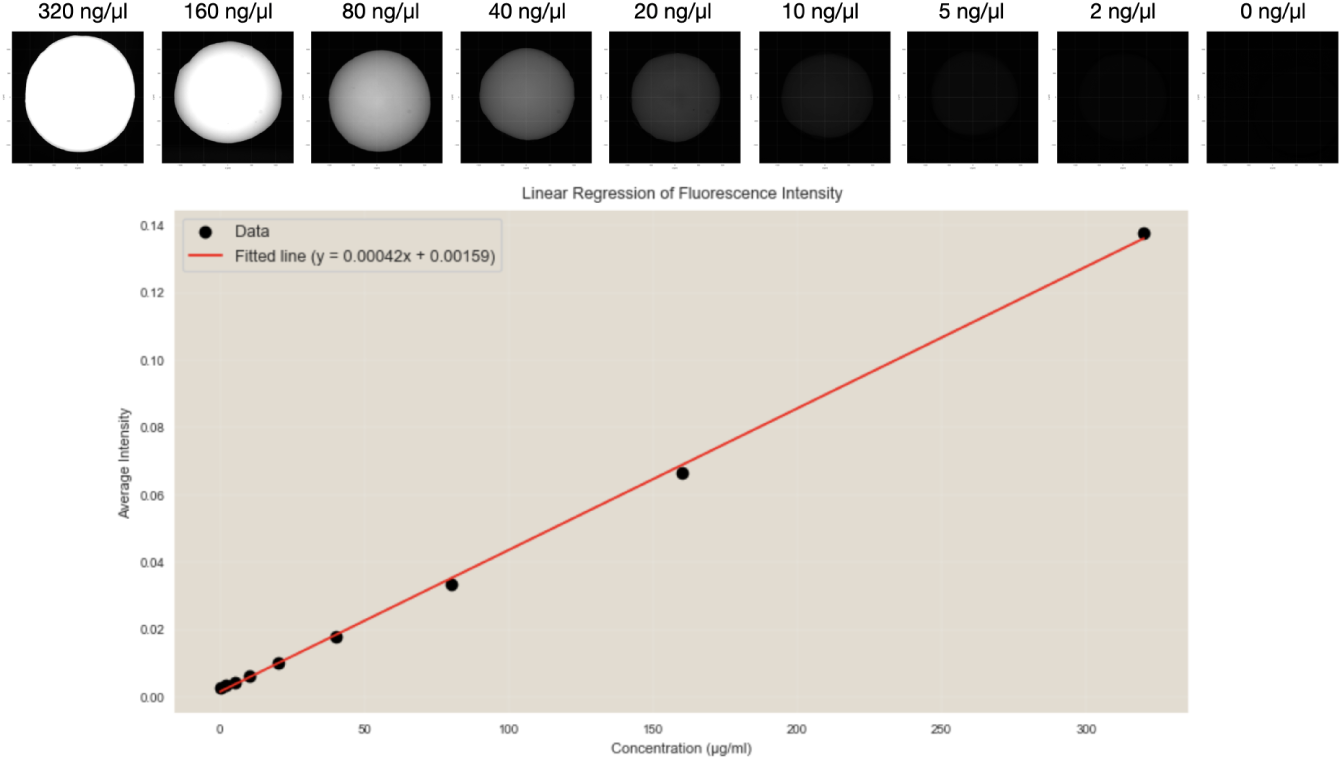
Calibration of mVenus fluorescence in ActiveDROPS droplets. Purified mVenus standards ranging from 0 to 320 ng/*µ*l were encapsulated and imaged using the droplet geometry and optical settings used for ActiveDROPS measurements (top). Mean droplet fluorescence intensity increased linearly with mVenus concentration over the measured range (bottom; (I = 0.00042C + 0.00159), with (C) in *µ*g/ml). The fitted calibration was used to estimate kinesin–mVenus concentration from droplet fluorescence.

**Fig. S3.**
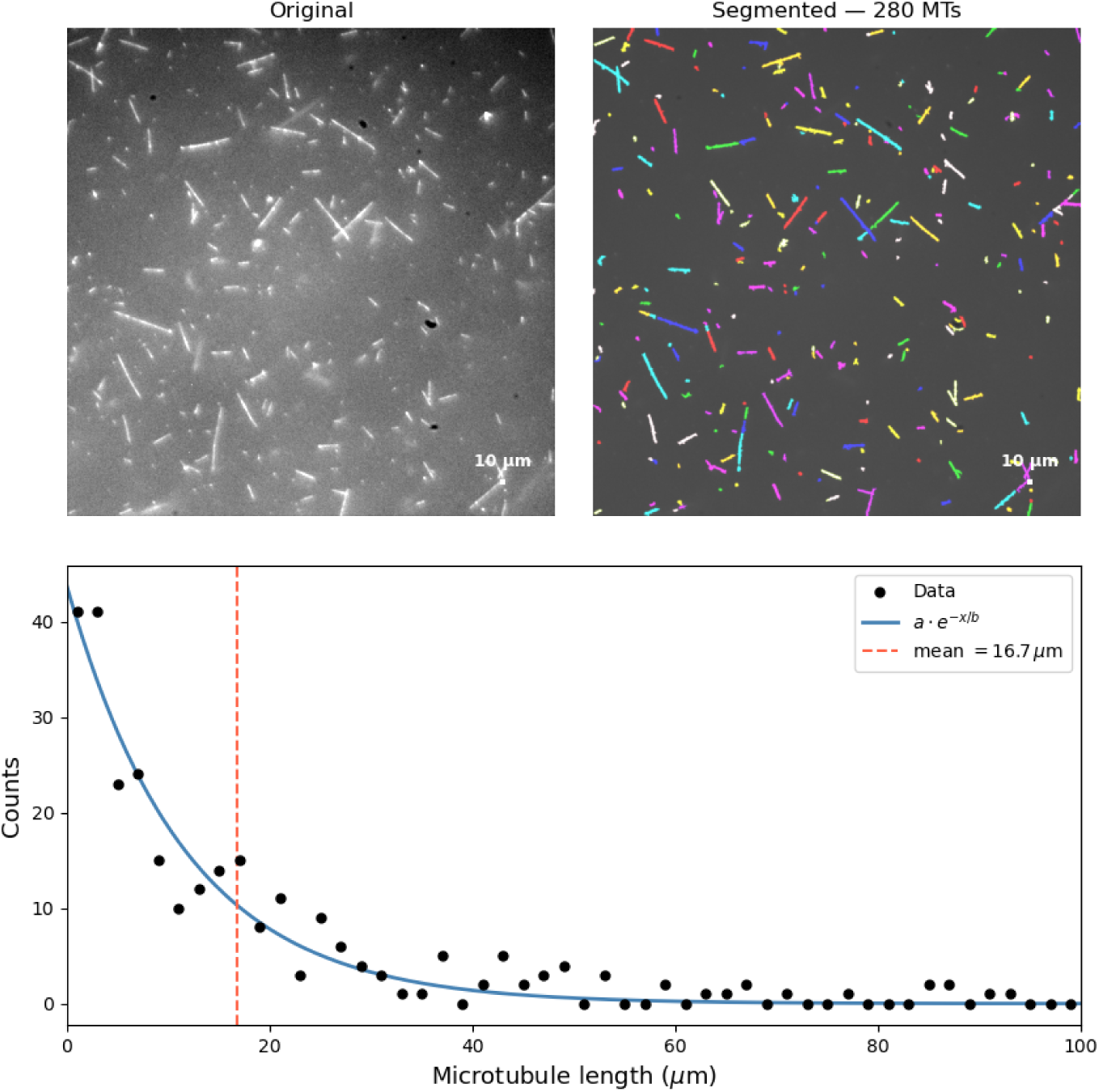
Length distribution of GMP-CPP-stabilized microtubules. (Top left) Representative fluorescence image of the stabilized microtubule preparation. Scale bar, 10 *µ*m. (Top right) Automated segmentation of the same field, containing 280 detected microtubules. (Bottom) Length histogram and exponential fit. The measured mean length is 16.7 *µ*m (red dashed line).

**Fig. S4.**
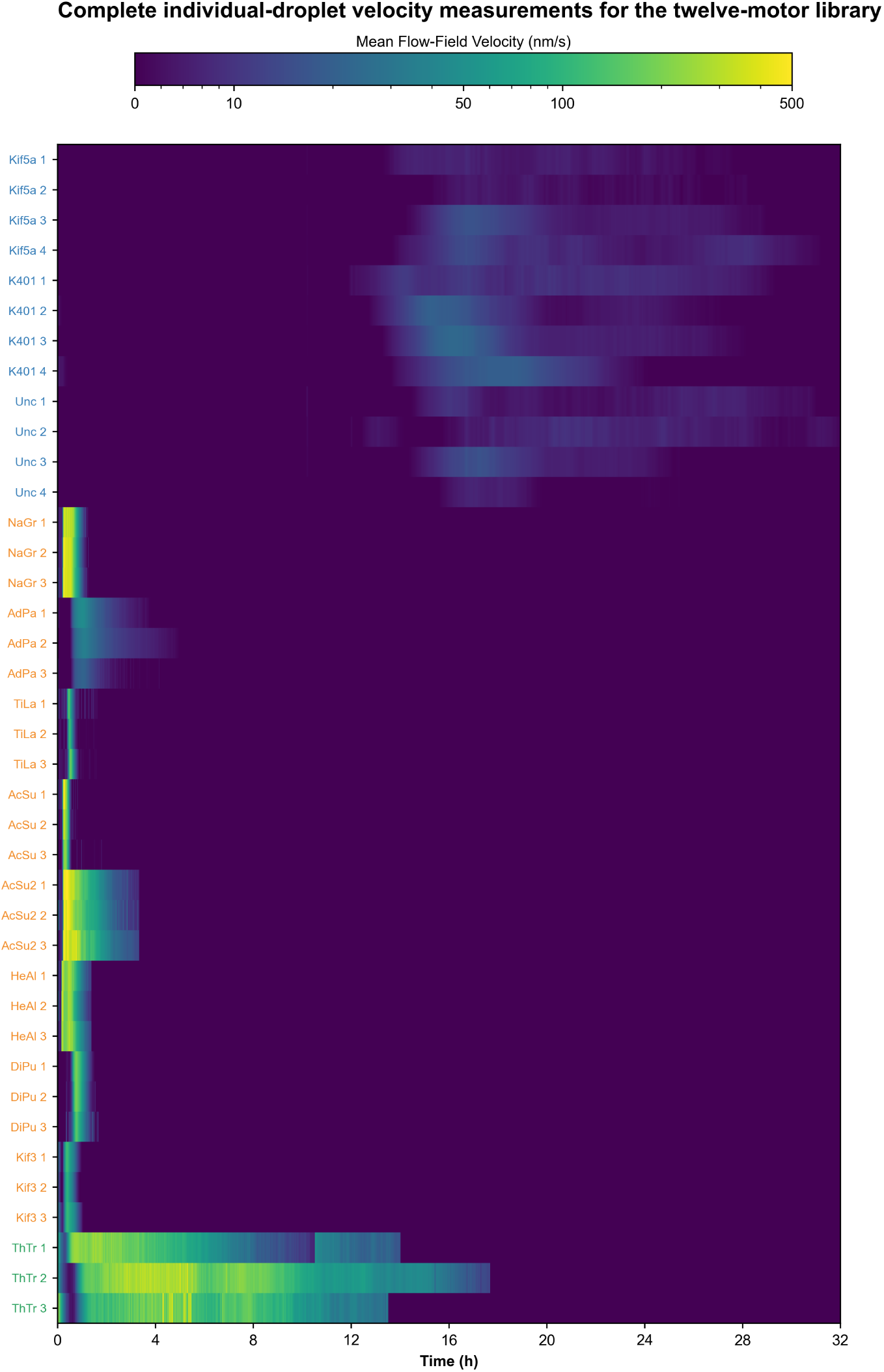
Complete individual-droplet velocity measurements for the twelve-motor library. Heatmap of mean flow-field velocity over absolute experimental time for all 39 independently prepared droplets used in the Figure 2 analysis. Rows are grouped and colored by the post hoc mechanical phenotypes Slow-Sustained, Fast-Burst, and Multiphase. Recorded inactive intervals and post-recording extensions are displayed at zero; for Fast-Burst recordings, extension follows the observed cessation of motion.

**Fig. S5.**
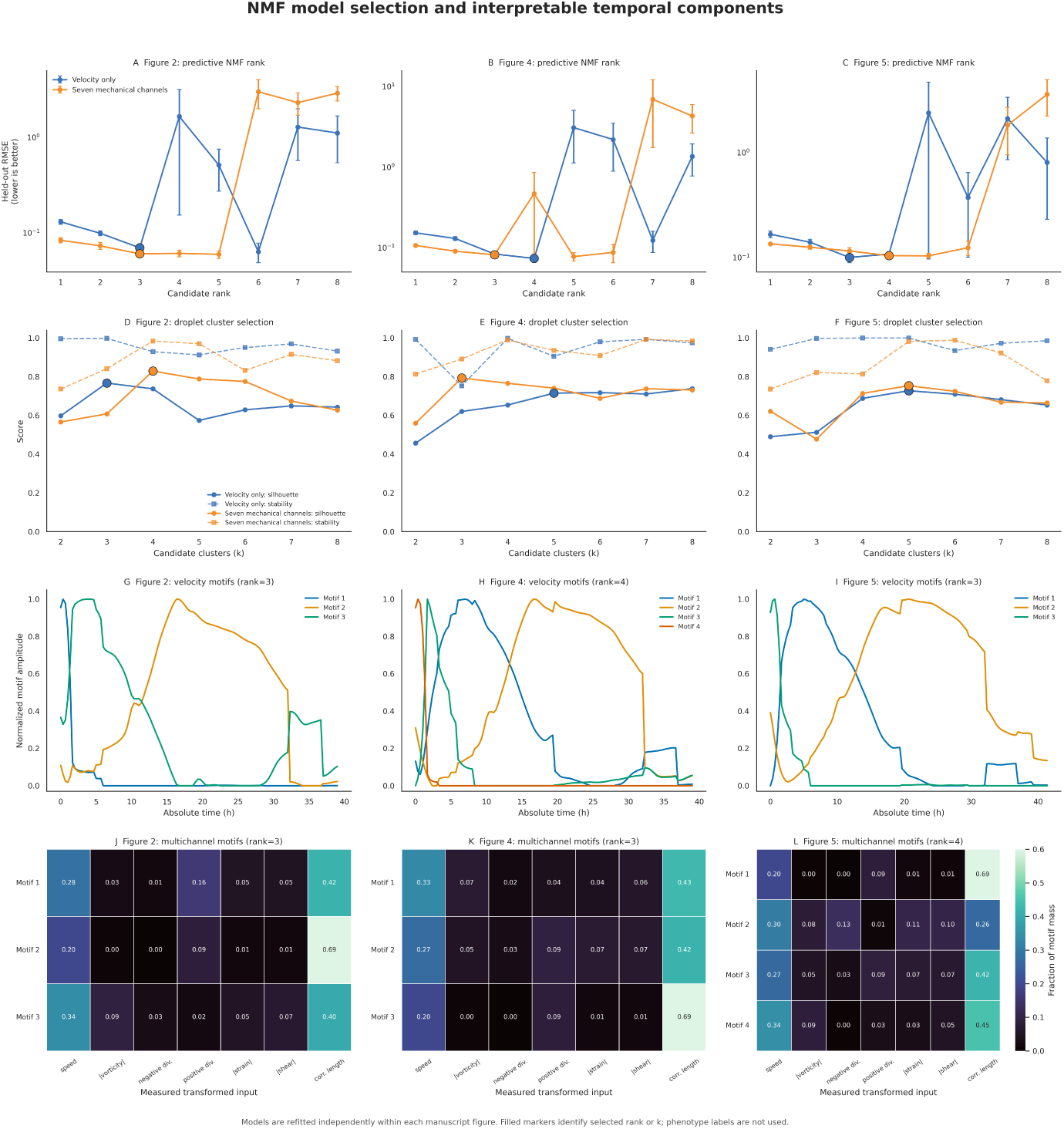
NMF model selection and interpretable temporal components. (A–C) Mean held-out reconstruction error for candidate NMF ranks in independently fitted Figure 2, Figure 4, and Figure 5 datasets. Error bars indicate the standard error across four contiguous held-out-time-block folds; filled markers identify the smallest rank within one standard error of the minimum error. Velocity-only NMF selected ranks 3, 4, and 3, whereas seven-channel NMF selected ranks 3, 3, and 4. (D–F) Silhouette coefficients (solid lines) and 80% droplet-subsampling stability measured by adjusted Rand index (dashed lines) for candidate cluster numbers in the complete standardized NMF-weight spaces. Filled markers identify the selected cluster numbers: (k=3), 5, and 5 for velocity-only NMF and (k=4), 3, and 5 for seven-channel NMF. (G–I) Temporal basis components learned from the zero-padded velocity histories. Components are normalized individually to display temporal shape; fitted droplet weights retain differences in component amplitude. (J–L) Relative contributions of velocity magnitude, absolute mean vorticity, negative and positive mean-divergence components, absolute mean strain, absolute mean shear, and correlation length to each multichannel NMF component. Phenotype, motor, construct, expression, and DNA-concentration labels were excluded from model selection and fitting.

**Fig. S6.**
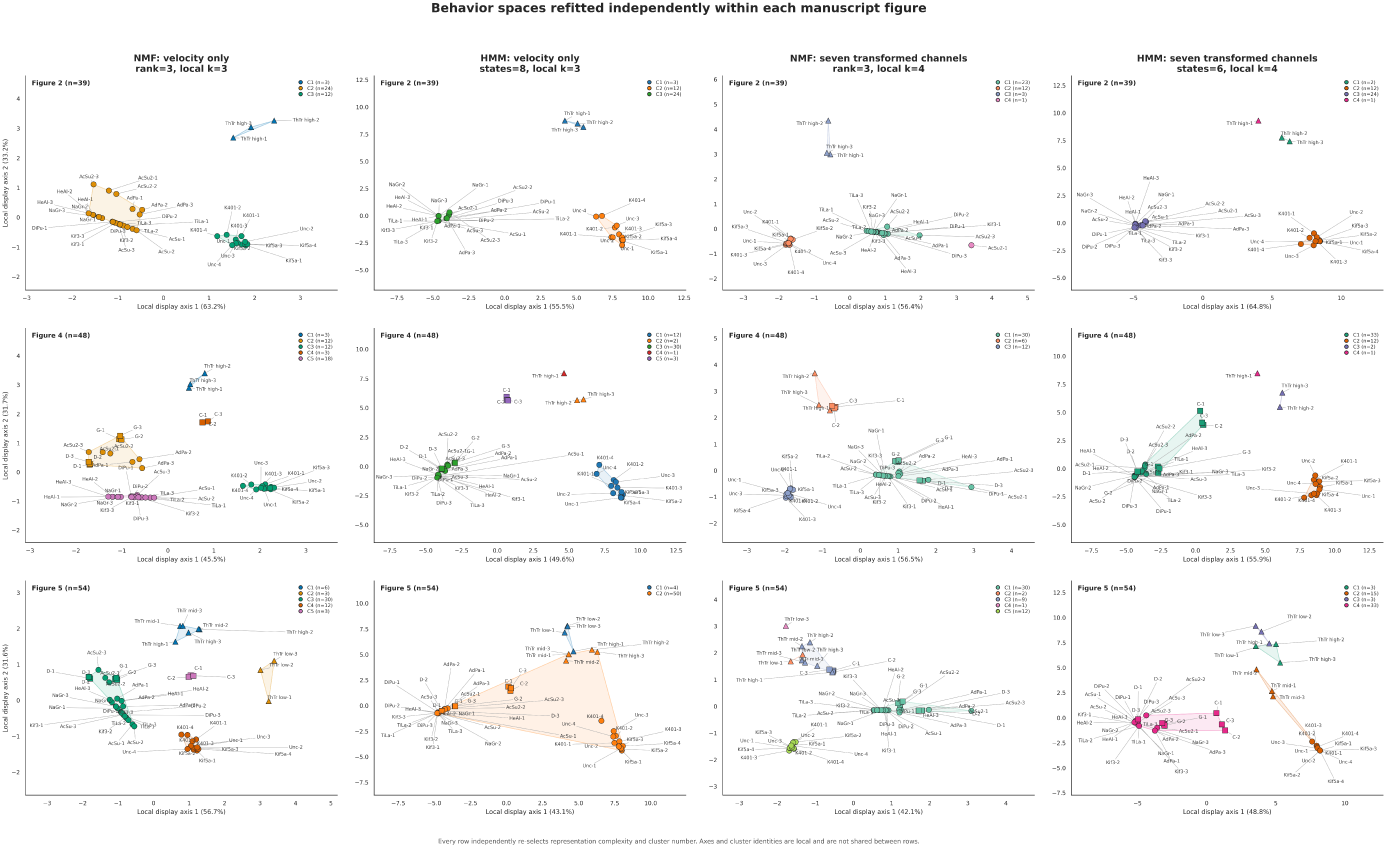
Comparison of independently refitted NMF and HMM behavior spaces. Behavior spaces for Figure 2 (39 droplets; top row), Figure 4 (48 droplets; middle row), and Figure 5 (54 droplets; bottom row) were independently constructed using, from left to right, velocity-only NMF, velocity-only HMM, seven-channel NMF, and seven-channel HMM. Each panel independently reselected representation complexity and droplet cluster number using only the droplets shown in that row. NMF clusters were calculated in the complete standardized weight spaces, whereas HMM clusters were calculated from pairwise Hamming distances between complete hidden-state paths. The two-dimensional PCA projections were used only for visualization and did not determine cluster membership. Colors denote label-free clusters and are local to each panel; corresponding colors in different panels do not imply equivalent cluster identities. Circles, squares, and triangles identify natural-motor, chimera, and ThTr expression-series droplets, respectively. Motor identity, construct identity, expression condition, DNA concentration, and phenotype labels were excluded from all fitting and model-selection steps.

**Fig. S7.**
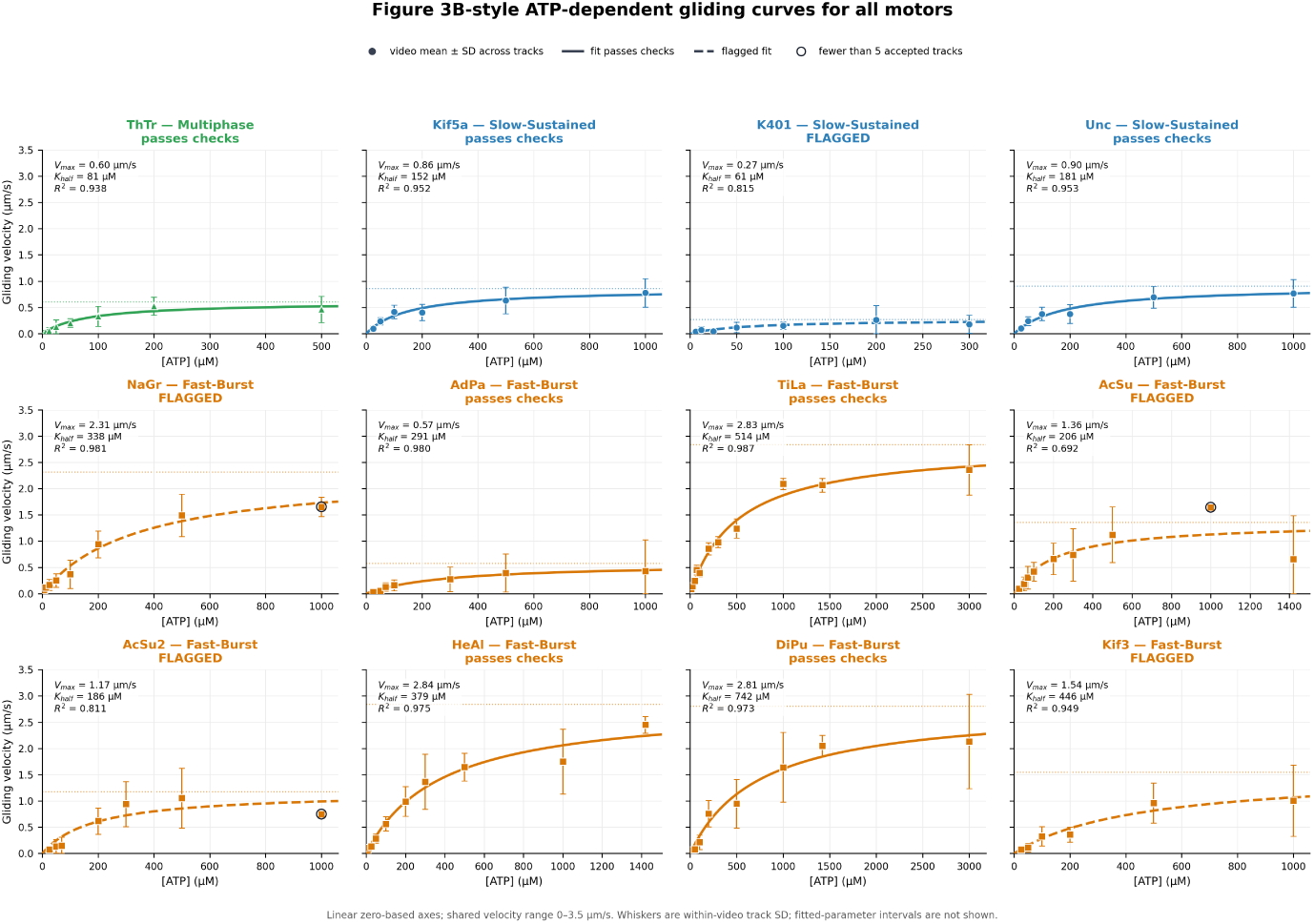
ATP-dependent gliding velocity across the twelve-motor library. Points show the mean *±* SD across accepted tracks within each video; curves show equal-weight descriptive Michaelis– Menten fits, with dotted lines indicating fitted *V*max. Dashed curves identify fits failing at least one predefined support criterion. ATP axes are linear and begin at zero.

**Fig. S8.**
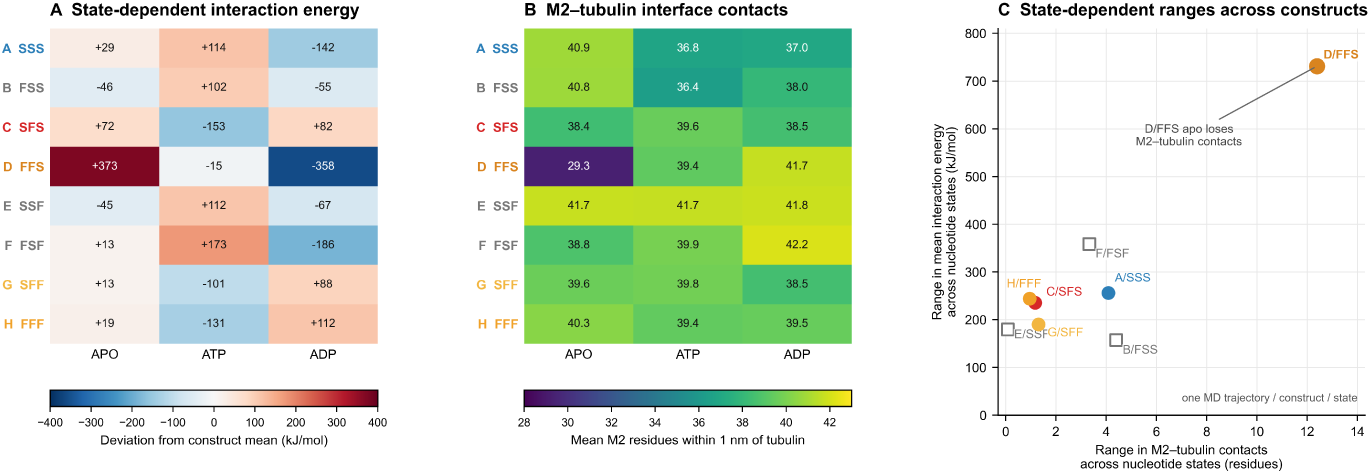
State-dependent motor–tubulin interaction energies and M2 interface contacts across the complete K401–Kif3 recombinant library. (A) Mean short-range Coulombic plus Lennard-Jones interaction energy for each construct in nucleotide-free (APO), ATP-bound, and ADP-bound simulations. Values were centered by subtracting each construct’s mean across the three nucleotide states; negative values indicate a more favorable interaction relative to that construct’s own mean. (B) Mean number of residues in the M2 crossover region located within 1 nm of tubulin during the corresponding simulations. (C) State-dependent variation in M2–tubulin contacts plotted against the variation in mean interaction energy, calculated as the maximum minus minimum value across the three nucleotide states. FFS showed the largest state-dependent range in both measurements, driven by reduced M2–tubulin contacts in the APO simulation. Filled circles denote constructs that generated detectable collective motion; open squares denote visually immobile constructs. Colors indicate Slow-Sustained (blue), Fast-Sustained (red), Fast-Burst (orange), or visually immobile (gray) behavior. Letters A–H correspond to the constructs in Figure 4; S and F indicate K401- and Kif3-derived regions, respectively. Each construct–state combination is represented by one 50-ns molecular-dynamics trajectory. These interaction energies are comparative direct-interaction measures, not binding free energies or experimental affinities.

**Fig. S9.**
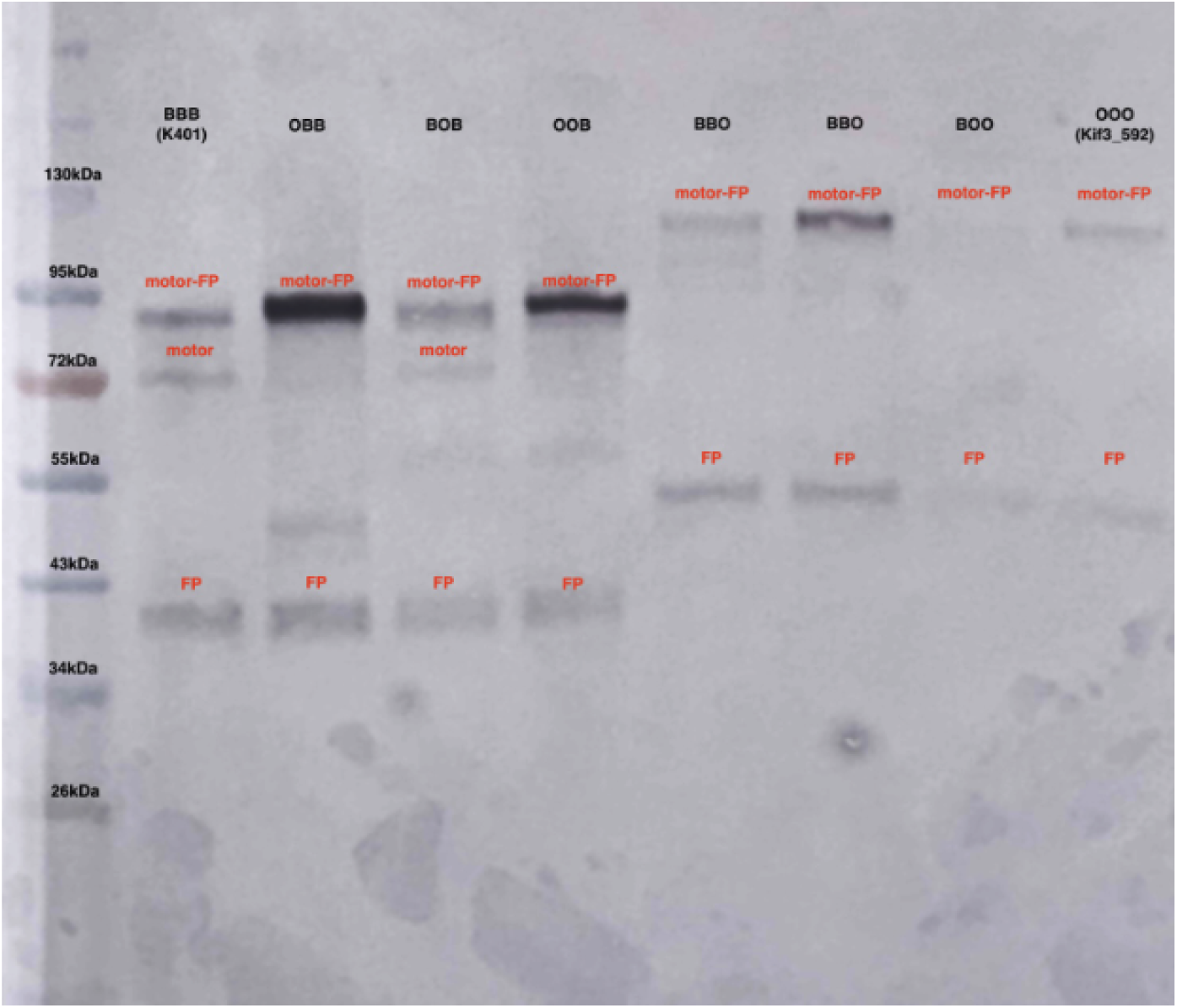
Western blot of TXTL reactions expressing all eight chimeric constructs. Proteins were probed with anti-mVenus antibody (anti-GFP clone B-2) targeting the C-terminal mVenus tag. Expected molecular weight for each chimera is indicated. All constructs produce a full-length band; a secondary band consistent with cleavage at the fluorescent-protein junction is visible in most lanes.

**Fig. S10.**
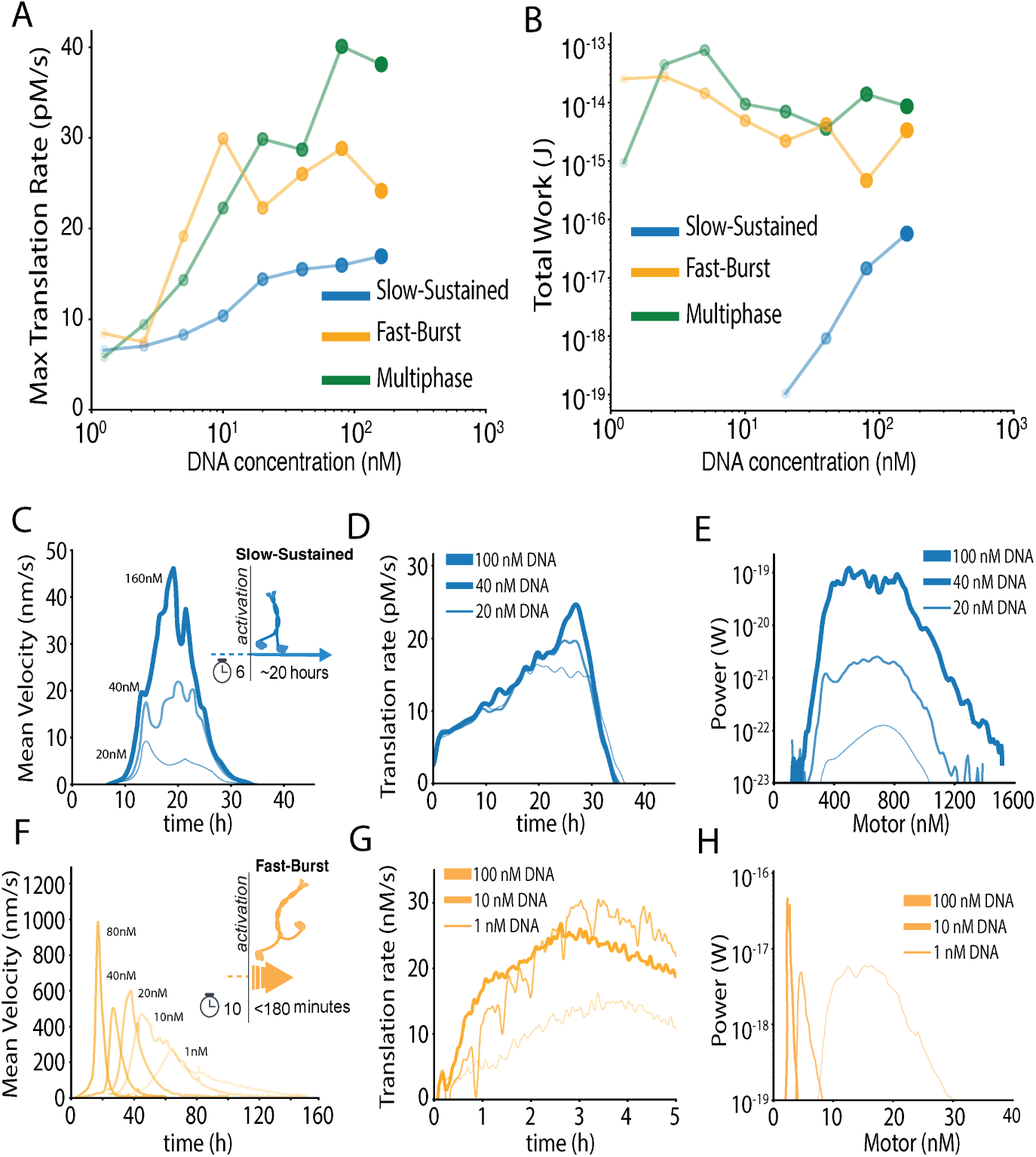
DNA input indirectly determines motor accumulation and mechanical output across self-organization phenotypes. (A) Maximum net kinesin-mVenus reporter-appearance rate as a function of input DNA concentration for Slow-Sustained, Fast-Burst, and Multiphase networks. (B) Total mechanical work generated as a function of DNA concentration. (C-E) Representative Slow-Sustained responses across the indicated DNA inputs. (C) Mean network velocity over time. (D) Net reporter-appearance rate over time. (E) Estimated mechanical work as a function of motor concentration. (F-H) Corresponding Fast-Burst responses across the indicated DNA inputs. (F) Mean network velocity over time. (G) Net reporter-appearance rate over time. (H) Estimated mechanical power as a function of motor concentration.

**Table S1.** Twelve kinesin-1 motors used in this study. Motors are referred to throughout the main text and Supplementary Information by the abbreviated names in the first column. Numerical suffixes indicate the C-terminal truncation residue of the construct relative to the native full-length sequence.

| Protein | Organism | Taxonomy | Accession ID |
| --- | --- | --- | --- |
| K401 | *Drosophila melanogaster* (animal) | Metazoa; Arthropoda; Diptera; Drosophilidae | P17210 |
| Kif3 | *Dictyostelium discoideum* (slime mold) | Amoebozoa; Dictyostelia; Dictyostelium | Q54UC9 |
| AcSu_575 | *Acytostelium subglobosum* (slime mold) | Amoebozoa; Dictyostelia; Acytostelium | XP_012759363 |
| AcSu2_520 | *Acytostelium subglobosum* (slime mold) | Amoebozoa; Dictyostelia; Acytostelium | BAN16545.10 |
| AdPa_408 | *Aduncisulcus paluster* (single-celled) | Metamonada; Fornicata; Aduncisulcus | GKT27654.1 |
| DiPu_513 | *Dictyostelium purpureum* (slime mold) | Amoebozoa; Dictyostelia; Dictyostelium | XP_003286323.1 |
| HeAl_513 | *Heterostelium album* (slime mold) | Amoebozoa; Dictyostelia; Heterostelium | XP_020435828.1 |
| Kif5a_500 | *Homo sapiens* (animal) | Metazoa; Chordata; Mammalia; Primates | NP_004975.2 |
| NaGr_392 | *Naegleria gruberi* (single-celled) | Heterolobosea; Naegleria | XP_002677830.1 |
| ThTr_655 | *Thecamonas trahens* (single-celled) | Apusozoa; Apusomonadida; Thecamonas | XP_013754985.1 |
| TiLa_514 | *Tieghemostelium lacteum* (slime mold) | Amoebozoa; Dictyostelia; Tieghemostelium | AGC69740.1 |
| Unc_554 | *Caenorhabditis elegans* (animal) | Metazoa; Nematoda; Rhabditida; Caenorhabditis | P34540 |

**Table S2.** Microtubule-gliding fit parameters for the twelve natural motors. Michaelis–Menten parameters were obtained from fits of gliding velocity as a function of ATP concentration. The apparent stepping rate is *V*max*/*(8 nm) and is used only as a velocity-derived estimate, not as a direct measurement of ATP turnover. The final column gives the ActiveDROPS phenotype assignment.

| **Motor** | $V_{\max}$<br>($\mu\text{m s}^{-1}$) | $K_{1/2}$<br>($\mu\text{M ATP}$) | **Apparent**<br>**step rate**<br>($\text{s}^{-1}$) | **Step rate/$K_{1/2}$**<br>($\mu\text{M}^{-1} \text{s}^{-1}$) | $R^2$ | $n_{\text{tracks}}$ | **Phenotype** |
| --- | --- | --- | --- | --- | --- | --- | --- |
| AcSu | 1.357 | 206 | 169.6 | 0.824 | 0.692 | 441 | Fast-Burst |
| AcSu2 | 1.175 | 186 | 146.9 | 0.790 | 0.811 | 311 | Fast-Burst |
| AdPa | 0.575 | 291 | 71.8 | 0.247 | 0.980 | 456 | Fast-Burst |
| DiPu | 2.808 | 742 | 351.0 | 0.473 | 0.973 | 358 | Fast-Burst |
| HeAl | 2.840 | 379 | 355.0 | 0.937 | 0.975 | 661 | Fast-Burst |
| Kif3 | 1.543 | 446 | 192.9 | 0.433 | 0.949 | 233 | Fast-Burst |
| NaGr | 2.312 | 338 | 289.1 | 0.855 | 0.981 | 263 | Fast-Burst |
| TiLa | 2.833 | 514 | 354.1 | 0.689 | 0.987 | 1984 | Fast-Burst |
| K401 | 0.267 | 61 | 33.4 | 0.547 | 0.815 | 195 | Slow-Sustained |
| Kif5a | 0.856 | 152 | 107.0 | 0.704 | 0.952 | 261 | Slow-Sustained |
| Unc | 0.904 | 181 | 113.0 | 0.626 | 0.953 | 302 | Slow-Sustained |
| ThTr | 0.604 | 81 | 75.5 | 0.929 | 0.938 | 189 | Multiphase |

**Table S3.** Motor–microtubule interaction energies across nucleotide states. Time-averaged nonbonded interaction energies (kJ mol*−*1) between the kinesin motor domain and the *αβ*-tubulin dimer, extracted from the final 30 ns of 50 ns GROMACS trajectories. Each value is averaged across three replicate simulations. Negative values indicate attractive interaction; more negative values indicate stronger interaction.

| **Motor** | $E_{\text{APO}}$ | $E_{\text{ATP}}$ | $E_{\text{ADP}}$ | **Class** |
| --- | --- | --- | --- | --- |
| AcSu | −1823.2 | −2160.6 | −2066.7 | Fast-Burst |
| NaGr | −2077.2 | −2007.8 | −1928.6 | Fast-Burst |
| TiLa | −2017.1 | −1893.3 | −2044.7 | Fast-Burst |
| HeAl | −1884.2 | −1888.8 | −2045.1 | Fast-Burst |
| DiPu | −1777.3 | −1866.9 | −1999.9 | Fast-Burst |
| Kif3 | −1849.7 | −1988.6 | −1777.4 | Fast-Burst |
| ThTr | −1821.0 | −1963.6 | −1852.5 | Multiphase |
| AdPa | −1811.9 | −1806.5 | −1849.7 | Fast-Burst |
| AcSu2 | −1662.0 | −1834.5 | −1853.7 | Fast-Burst |
| Kif5a | −1699.9 | −1742.8 | −1809.8 | Slow-Sustained |
| K401 | −1540.6 | −1551.3 | −1754.4 | Slow-Sustained |
| Unc | −1489.0 | −1621.6 | −1505.9 | Slow-Sustained |

**Table S4.** Kinesin motor-domain subdomains: structural elements, functional roles, and mechanochemical contributions.

| Subdomain (module) | Residues | Role in ATP cycle | Role in MT interaction | Reference |
| --- | --- | --- | --- | --- |
| P-loop (M1) | GxxxxGKT, Y96 | Binds ATP and $Mg^{2+}$; stabilizes nucleotide for hydrolysis | Indirect: stabilizes ATP pocket | Song et al. |
| $\alpha 0$ helix (M1) | — | Positions L5 and Sw-I for ATP entry | Indirect via ATP-pocket stabilization | Hwang et al. |
| Loop 5 (M1) | T101 | Positions adenosine ring; aligns ATP for hydrolysis | Indirect | Song et al. |
| Central β-sheet (M1) | core strands | Maintains pocket geometry | Indirect; force transmission | Song et al. |
| Switch-I (M2) | R203, SSR motif | γ-phosphate sensor; closes on ATP | Stabilizes closed state; raises MT affinity | Hwang et al.; Song et al. |
| Switch-II (M2) | R234–E270 salt bridge, E236 | Coordinates hydrolysis with Sw-I; aids Pi release | Interacts with L11, L12; reinforces MT attachment | Hwang et al.; Cao et al. |

**Table S4 Kinesin motor-domain subdomains: structural elements, functional roles, and mechanochemical contributions (continued).**
| Subdomain (module) | Residues | Role in ATP cycle | Role in MT interaction | Reference |
| --- | --- | --- | --- | --- |
| Sw-I/Sw-II salt bridges (M2) | R203–E236, R207–E240 | Stabilize ATP-bound state; lower hydrolysis barrier | Indirect via state stabilization | Song et al. |
| Loop 11 (M2) | G243 (NcKin); basic residues | Not directly catalytic | Primary MT contact; gates processivity | Cao et al.; Song et al. |
| $\alpha 4$ helix (M2) | basic residues | Conformation tracks nucleotide state | Anchors L11 on MT; key for power stroke | Song et al. |
| $\alpha 5$ helix and L12 (M2) | basic residues | Indirect | Reinforces MT attachment during stress | Hwang et al. |
| Neck linker (M3) | proximal residues | ATP-driven docking drives forward step | Coordinates head positioning on MT | Hwang et al.; Song et al. |

